# PTBP2 promotes cell survival and autophagy in Chronic Myeloid Leukemia by stabilizing BNIP3

**DOI:** 10.1101/2024.01.26.577177

**Authors:** Bibhudev Barik, Shristi Lama, IS Sajitha, Sayantan Chanda, Sonali Mohapatra, Sutapa Biswas, Ghanashyam Biswas, Soumen Chakraborty

## Abstract

Polypyrimidine tract binding protein 2 (PTBP2) regulates alternative splicing in neuronal, muscle, and Sertoli cells. PTBP2 and its paralog, PTBP1, which plays a role in B-cell development, was found to be expressed aberrantly in myeloid leukemia. Genetic ablation of Ptbp2 in the cells resulted in decreased cellular proliferation and repopulating ability, decreased reactive oxygen species (ROS), and altered mitochondrial morphology. The sensitivity of CML cells to imatinib increased after the knockout of Ptbp2. RNA immunoprecipitation followed by sequencing (RIP-seq) and functional assays confirmed that PTBP2 binds to Bcl-2 Interacting Protein 3 (Bnip3)-3’UTR and stabilizes its expression. Our study also suggests that PTBP2 promotes autophagy, as evidenced by the low levels of LC3-II expression in Ptbp2-knockout cells treated with Bafilomycin A1. This effect was restored upon overexpression of Bnip3 in the knockout cells. Notably, when KCL22-NTC cells were subcutaneously injected into the flanks of mice, they gave rise to malignant tumors, unlike Ptbp2-KO-KCL22 cells. This underscores the role of PTBP2 in promoting cell proliferation and tumor formation while enhancing autophagy through Bnip3, thereby supporting the role of PTBP2 as an oncogene in CML.

## Introduction

Chronic Myeloid Leukemia (CML) is a type of blood cancer that arises due to a genetic translocation between chromosomes 9 and 22, leading to the expression of a constitutively active tyrosine kinase oncogene called Bcr-Abl1 (**Osman and Deininger, 2021**). The resultant oncoprotein triggers pro-survival signaling pathways in CML cells, providing them with a proliferative advantage and resistance to apoptosis. While tyrosine kinase inhibitors (TKIs) have significantly improved the survival rate of patients with chronic phase CML, the survival rate is much lower for those in the blast phase (**Hochhaus et al., 2017; Jain et al., 2017**). Additionally, most patients require ongoing maintenance therapy for the rest of their lives, as current treatments only suppress the growth of cancer cells. Various BCR-ABL1-dependent and -independent pathways, such as kinase domain mutations, Bcr-Abl1 amplification, and aberrant activation of the PI3K and RAS/MAPK signaling pathways, increase the risk of progression and predict poorer response to TKIs (**Melo and Barnes, 2007; Diamond and Melo, 2011; Wagle et al., 2016**). Therefore, identifying the factors responsible for disease progression is crucial to enhance the treatment of CML, particularly in advanced cases. Recent studies have highlighted the crucial role of RNA-binding proteins (RBPs) in cancer by regulating mRNA expression. The polypyrimidine tract-binding protein (PTBP) is one such RBP that plays a critical role in regulating mRNA stability, protein expression, and exon exclusion, thereby influencing cellular growth and development (**Sawicka et al., 2008**). PTBP2, a subtype of PTBP, exhibits differential tissue expression and is correlated with several types of cancer, including glioblastoma, osteosarcoma, and colorectal cancer (**Yang et al., 2014; Ji et al., 2014; Sun et al., 2019; Kim et al., 2021**). Autophagy, a cellular process that maintains cellular balance and prevents the accumulation of impaired proteins and organelles, has also been implicated in tumorigenesis (**Frankel et al., 2017; Abildgaard et al., 2020; Glick, et al., 2010**). Its role in CML is complex, with studies suggesting that it can contribute to both the survival and proliferation of CML cells as well as the development of drug resistance (**Helgason, et al., 2013; Mishima et al., 2008; Bellodi et al., 2009**).

This study investigated the role of PTBP2 in regulating CML pathology and found that it promotes cell proliferation through oxidative phosphorylation (OXPHOS) supported by mitochondrial fusion. Additionally, PTBP2 promotes autophagy by binding and stabilizing Bnip3, which has been implicated in cancer cell subpopulations and can be a potential therapeutic target. Mice injected with Ptbp2 KO cells exhibited significantly smaller tumors and reduced proliferation and autophagy markers, supporting the role of PTBP2 as an oncogene in CML.

## Results

### Establishment and characterization of PTBP2 knockout CML and AML cells

In this study, we examined PTBP2 expression in CML and AML cell lines. The expression of PTBP2 was nearly identical in the four CML cell lines, KCL22, K562, KU812, and KYO1 (**Fig. 1A**). In one CML cell line, LAMA84, PTBP2 expression was found to be reduced (**Fig. 1A**). TF1 (human erythroleukemia), HEL (human erythroid leukemia), and F36P (MDS-RAEB) showed varying levels of PTBP2 expression, whereas HNT34 (BCR-ABL1+ human Acute Myeloid Leukemia) cells showed almost no PTBP2 expression among the four AML cell lines (**Fig. 1A**).

**Figure 1.**
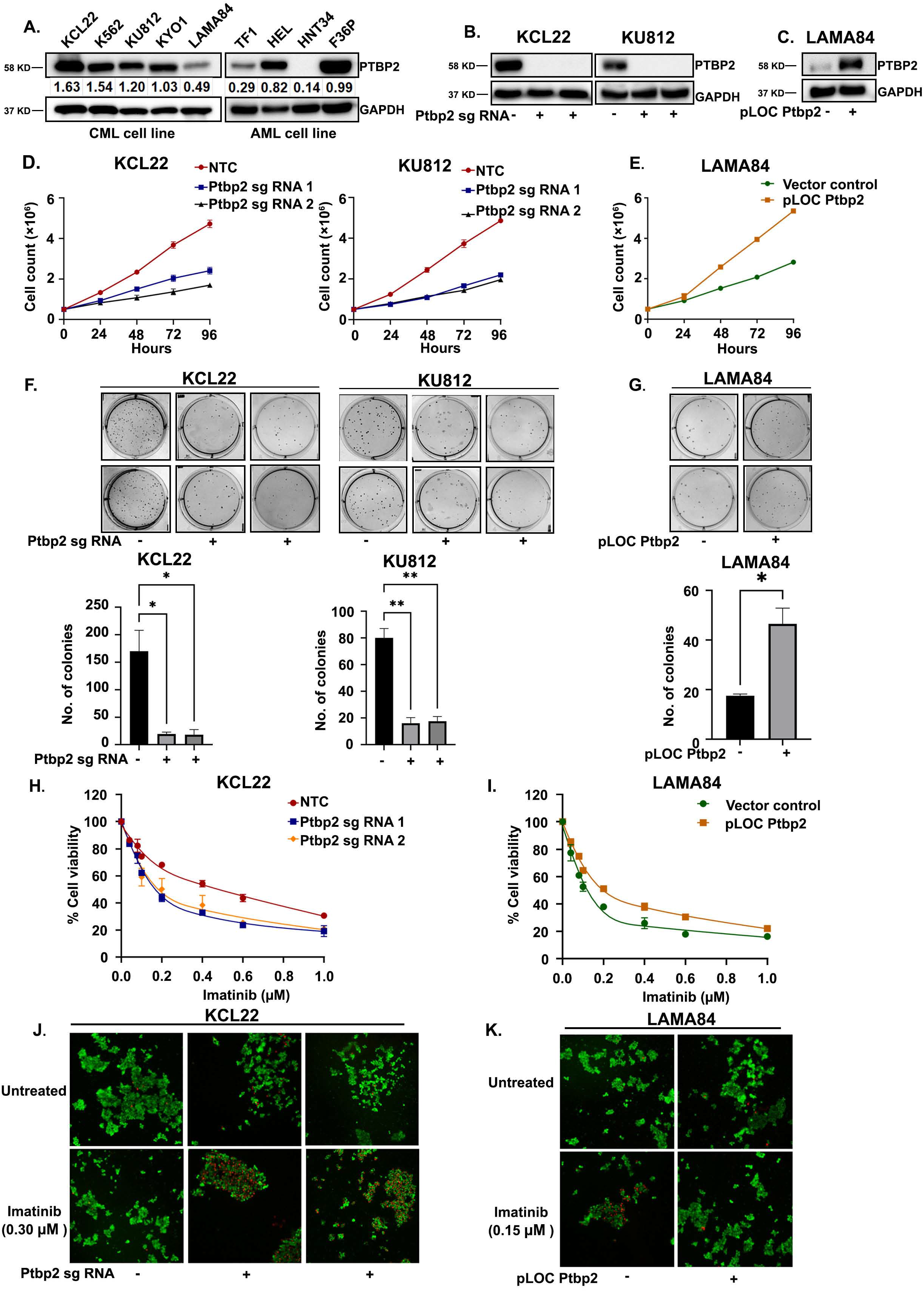
PTBP2 protein expression in different cell types and phenotypic assessment. **A**. PTBP2 protein expression in CML cell lines (KCL22, K562, KU812, KYO1, and LAMA84) (lanes 1-5) & AML cell lines (TF1, HEL, HNT34, and F36P) (lanes 6-9). An equal amount of protein (30μg) was loaded, and GAPDH western blotting was used as an internal loading control. Densitometry quantification is shown in the figure. **B**. Western blot analysis of KCL22-NTC and Ptbp2-KO-KCL22, KU812-NTC, and Ptbp2-KO-KU812 cell lysates probed with the indicated antibody. The GAPDH antibody was used as an internal loading control. **C**. Western blot of PTBP2 transduced and stably selected LAMA84 (Ptbp2 OE LAMA84), and vector control cell lysate was probed with the indicated antibody. The GAPDH antibody was used as an internal loading control. **D**. The proliferation rates of NTC KCL22-NTC, KU812-NTC, and Ptbp2-KO-KCL22, Ptbp2-KO-KU812 cells were evaluated using the trypan blue exclusion test for cell viability. Proliferation rates at 24, 48, 72, and 96h are represented in the line graph. Each point represents the mean and standard deviation of independent triplicates. **E**. LAMA84 vector control and Ptbp2-OE-LAMA84 cell proliferation rate were assessed using a trypan blue dye exclusion test for cell viability. Proliferation rates at 24, 48, 72, and 96h are represented in the line graph. Each point represents the mean and standard deviation of independent triplicates. **F**. Soft agar colony formation assay was performed using KCL22-NTC, KU812-NTC, Ptbp2-KO-KCL22, and Ptbp2-KO-KU812 cells. Colony numbers are represented in the bar graph below the respective Fig. The bar diagram shows the mean value and corresponding standard deviation for the represented data: *n*=2 \**p*<□0.05 and \*\**p*<□0.01. **G**. Soft agar colony formation assay was performed using LAMA84 vector control and Ptbp2-OE-LAMA84 cells. Colony numbers are represented in the bar graph below the Figure. The bar diagram shows the mean value and corresponding standard deviation for the represented data; *n*=2 \**p*<□0.05. **H**. Representative graph of % cell viability in KCL22-NTC and Ptbp2-KO-KCL22 cells by MTT assay after treatment with variable imatinib doses. **I**. Representative graph of % cell viability in LAMA84 vector control and Ptbp2-OE-LAMA84 cells treated with indicated doses of imatinib. **J**. Representative images of imatinib treated and untreated KCL22-NTC, Ptbp2-KO-KCL22 cells from Cell Insight CX7 High Content Screening (HCS) Platform. **K**. Representative images of imatinib treated and untreated LAMA84 vector control & Ptbp2-OE-LAMA84 cells from Cell Insight CX7 High Content Screening (HCS) Platform.

Next, we assessed cell proliferation and colony formation ability in soft agar by ablating Ptbp2 in CML and AML cells. Previously, we reported that the knockdown of Ptbp2 facilitated the reduction in cellular growth and increased apoptosis **(Nandagopalan et al., 2019**). Herein, Ptbp2 knockout (KO) was achieved using two distinct gRNAs, as outlined in the Materials and Methods, and several single-cell knockout clones for each cell line were generated and propagated. Using RT-qPCR and western blotting, we confirmed the extent of Ptbp2 knockout in KCL22, KU812, and TF1 cells (**Fig. 1B and Supplementary Fig. 1A and B**) with respect to the non-targeting control (NTC) cells. Additionally, Ptbp2 overexpression (OE) was confirmed in the LAMA84 cells (**Fig. 1C and Supplementary Fig. 1C**). To determine the proliferation rate of the cells, we conducted a trypan blue dye exclusion test on both NTC and Ptbp2-KO-KCL22, KU812, TF1 cells. As previously noted, the growth rate was decreased in the Ptbp2-KO-KCL22, KU812, and TF1 clones (**Fig. 1D and Supplementary Fig. 1D**). An increase in the proliferation rate was observed upon overexpression of Ptbp2 in LAMA84 cells (**Fig. 1E**). Furthermore, the colony count was also found to be lower in the Ptbp2-KO-KCL22, KU812, and TF1 cells than in the NTC cells (**Fig. 1F and Supplementary Fig. 1E**). PTBP2 overexpressed LAMA84 cells showed a higher number of colonies than vector control cells (**Fig. 1G**). The bar graphs shown in **Fig. 1F** and **1G** and **Supplementary Fig. 1E** indicate a significant change in the colony count.

Although imatinib has revolutionized CML treatment, in cases where patients do not respond well or develop resistance, second-generation TKIs, such as dasatinib and nilotinib, are used. From MTT cell viability assay, a significant decrease in the IC50 value was observed upon imatinib treatment in Ptbp2-KO-KCL22 cells compared to the NTC cells **(Fig. 1H)**; however, treatment with dasatinib and nilotinib showed no significant changes (**Supplementary Fig. 1F**). After 48h of drug exposure, the IC50 values were 0.480 and 0.167μM in KCL22-NTC and Ptbp2-KO-KCL22 cells, respectively (**Fig. 1H**). In addition, overexpression of Ptbp2 in the LAMA84 cells increased the IC50 value to 0.197μM. It was 0.121μM for the vector control **(Fig. 1I)**. Again, upon imatinib treatment, more dead cells (Red) were observed in Ptbp2-KO-KCL22 and vector-LAMA84 cells with respect to the KCL22-NTC, and PTBP2-OE-LAMA84 cells, respectively (**Fig. 1J and 1K**). Our findings indicate that PTBP2 promotes viability in CML cells and protects cells from imatinib-mediated cell death.

### PTBP2 targets, stabilizes and regulates Bnip3 in CML cells

PTBP2 functions as an RNA-binding protein and its four RNA-binding domains regulate the pre-mRNA splicing, translation, and stability of target mRNAs. Therefore, we performed RNA immunoprecipitation and sequencing (RIP-seq) in the KCL22 cell line, which endogenously expresses high levels of PTBP2. First, the complex containing PTBP2 protein and its associated RNA was immunoprecipitated with a PTBP2-specific antibody (**Fig. 2A**), and subsequently, RNA sequencing was performed to profile the PTBP2-bound transcripts. Peak calling with RIPSeeker was performed on each IP alignment file, which produced a list of peaks and their corresponding candidate transcripts. Bound transcripts showing the maximum read counts were considered. Combining three biological repeats, we found that 24 mRNAs were enriched in the PTBP2 RNA-IP samples compared to the control IgG-IP samples (**Fig. 2B and 2C**). Gene Ontology Biological Process (GOBP) analysis of these mRNAs was performed to explore the signaling pathways that PTBP2 might control in CML cells. The Cluster Profiler Bioconductor package was used for gene ontology enrichment of UP- and DOWN-regulated genes for all comparisons with a p-value cutoff of less than or equal to 0.05. The top 10 enriched GO terms were visualized as dot plots (**Supplementary Fig. 2A**). GO analysis identified significant pathways that regulate transmembrane transport, transmembrane ion transport, and membrane potential, which consists of mitochondrial membrane potential. The mRNA candidates observed in this group were Nampt, Bnip3, and Cacnb2. The binding efficiency between the mRNAs and PTBP2 was validated by independent RNA-IP experiments followed by RT-qPCR, confirming the association between PTBP2 and its targets. Of the 24 targets, strong binding efficiency was detected in 11 (**Fig. 2D**). The remaining 13 exhibited no notable changes. Nampt, followed by Bnip3, showed the highest binding efficiency. The expression levels of 24 targets were compared by RT-qPCR between KCL22-NTC and Ptbp2-KO-KCL22 cells and are represented in a heat map (**Fig. 2E**). Although the binding affinity to Nampt was the highest, no corresponding change in NAMPT protein expression was detected between KCL22-NTC and Ptbp2-KO-KCL22 cells (**Fig. not shown**). Thus, the 2^nd^ top target, Bnip3 (Bcl-2 Interacting Protein 3), was considered, and its role was further explored in the pathology of CML.

**Figure 2.**
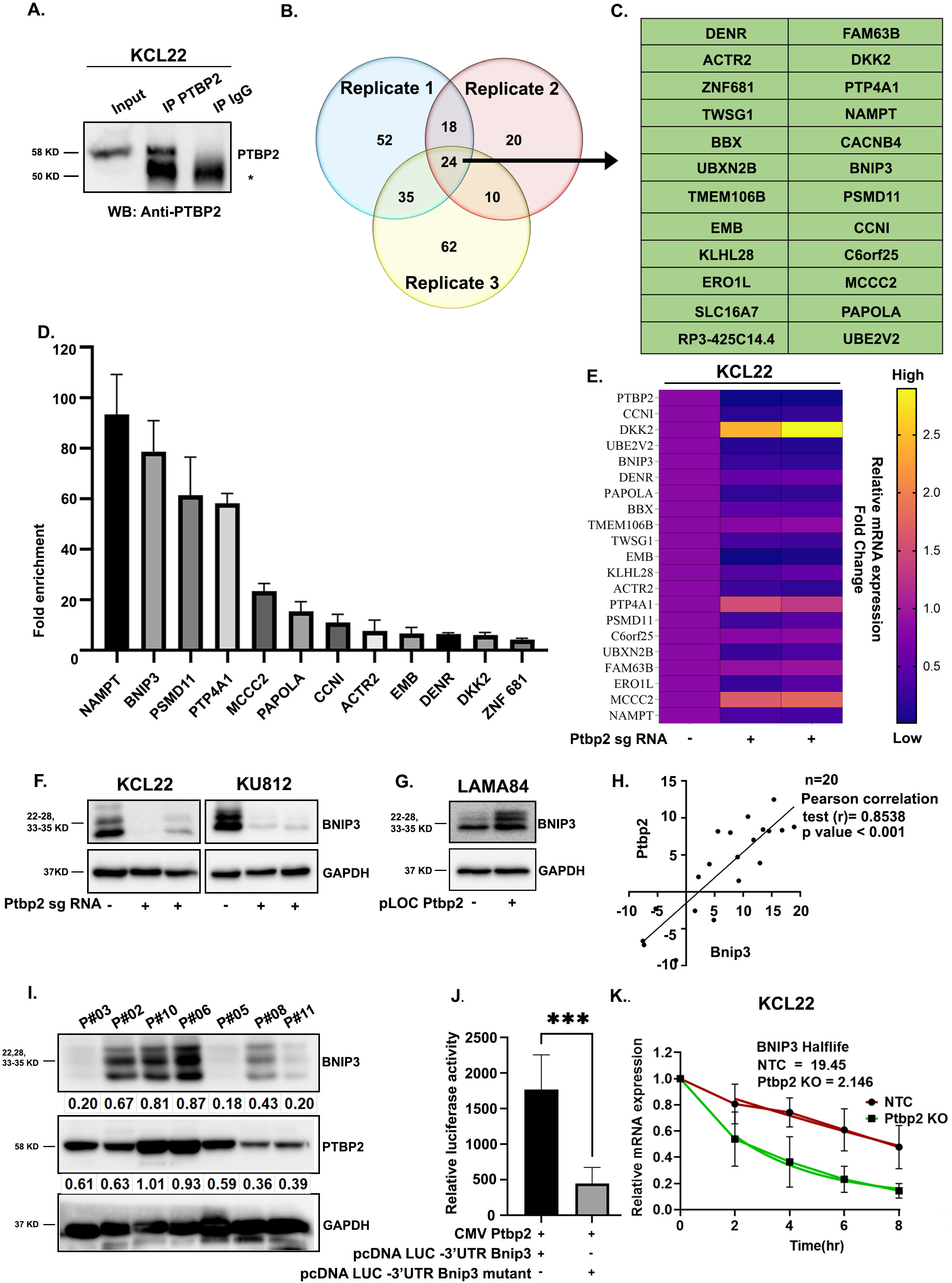
PTBP2 binds and regulates BNIP3. **A**. Immunoprecipitation of PTBP2 was performed using a PTBP2 antibody; 10% of the whole cell lysate was used as input lysate (lane 1), lysate immunoprecipitated with PTBP2 antibody (lane 2), lysate immunoprecipitated with IgG as a negative control (lane 3). The blot was probed with a PTBP2 antibody. “*” represents the IgG. **B**. Each circle represents the targets obtained from the single RIP-sequence experiment. Three repeats are presented in the Venn diagram. **C**. The common PTBP2-bound targets are represented in the table. **D**. RT-qPCR analysis of the PTBP2 bound target transcripts showing a fold enrichment with IgG control. **E**. Heat map showing a fold change in relative mRNA expression of the targets in KCL22-NTC and Ptbp2-KO-KCL22 cells. **F**. Whole-cell lysates of KCL22-NTC, KU812-NTC, and Ptbp2-KO-KCL22 and Ptbp2-KO-KU812 cells were collected, immunoblotting was performed with BNIP3 antibody, and GAPDH was used as an internal loading control. **G**. Whole-cell lysate of LAMA84 vector control and Ptbp2-OE-LAMA84 cells were collected, immunoblotting was performed with BNIP3 antibody, and GAPDH was used as an internal loading control. **H**. Ptbp2 and Bnip3 mRNA correlation in CML patient sample (n=20). **I**. Immunoblotting of some patient samples probed with BNIP3, PTBP2, and GAPDH antibodies. Densitometric quantification is shown in the figure. **J**. 3’UTR binding site in Bnip3 & the mutated site was cloned in the luciferase vector, and luciferase assay was performed by transfecting the clone along with wild-type PTBP2 in HEK293T cells. Renilla luciferase was used as an internal control. After 24h, luciferase activity was measured. Data presented are the fold change relative to empty vector-transfected cells. The data was analyzed and represented by a bar diagram. \*\*\**p*<□0.001. **K**. Half□life of the BNIP3 expression is represented for KCL22-NTC and Ptbp2-KO-KCL22 cells.

The role of the BH3-only protein BNIP3 in cancer is controversial. Increased BNIP3 levels in cancer patients have been linked to good as well as poor prognosis, as BNIP3 contributes to both pro-cell death and pro-survival signals **(Zhu et al., 2013; Singh et al., 2018)**. RT-qPCR and western blot results illustrated that the expression of BNIP3 in KCL22, KU812, and TF1 NTC cells was higher than that in Ptbp2-KO cells (**Fig. 2F and Supplementary Fig. 2B**). An increase in the expression of Bnip3 was noted when PTBP2 was overexpressed in LAMA84 cells (**Fig. 2G and Supplementary Fig. 2C**). Analysis of a publicly available dataset GSE4170, showed that as the disease progressed, there was a gradual increase in the expression of both the genes (**Supplementary Fig. 2D**) (**Radich et al., 2006**). An independent set of CML samples (n=20) also showed a positive Pearson correlation (r) of 0.8538 (*p*<0.001) between Ptbp2 and Bnip3 expression (**Fig. 2H**). Western blot analysis of some patient samples showed the same correlation (**Fig. 2I**). Thus, our study revealed that Ptbp2 strongly correlates with Bnip3 in CML.

As PTBP2 is an RNA-binding protein, and Bnip3 is one of the most enriched targets in RIP sequencing as well as RT-qPCR validation, we wanted to look into the binding site(s) of PTBP2 on Bnip3 mRNA, for which we scanned Bnip3 mRNA using beRBP (https://bioinfo.vanderbilt.edu/beRBP/predict.html). Only one specific PTBP2 binding site (CUUUUCU) was observed in the 3’-UTR, which had a binding score of 0.362 (**Supplementary Fig. 2E**). As we observed downregulation of Bnip3 in the absence of PTBP2, we speculate that PTBP2 binds to the 3’UTR of Bnip3 to stabilize it. Next, to confirm the binding of PTBP2 to Bnip3, we cloned the predicted site into a luciferase vector. Transfection with recombinant Ptbp2 (kindly gifted by Dr. Miriam Llorian, University of Cambridge, Cambridge, UK) increased the stability of the relative luciferase reporter count compared to the empty vector control. Modification of the binding site (CAAAACA) using site-directed mutagenesis decreased the relative luciferase reporter count, providing evidence that PTBP2 binds to and regulates the stability of Bnip3 (**Fig. 2J and Supplementary Fig. 2E**). The analysis of BNIP3 mRNA half-life as a measure of mRNA stability was investigated after transcriptional inhibition with Actinomycin D. We observed a noticeable impact on the stability of BNIP3 when PTBP2 was knocked out from the cells (**Fig. 2K**). Thus, our findings indicate that PTBP2 binds and functions through BNIP3.

### PTBP2-mediated oxidative phosphorylation supported by mitochondrial fusion increases cell proliferation

As cell proliferation is directly linked to cellular energy metabolism, we investigated the mitochondrial function (i.e., OCR) and substrate-level phosphorylation via glycolysis (i.e., ECAR) of KCL22-NTC and Ptbp2-KO-KCL22 cells and LAMA84 vector control and Ptbp2-OE-LAMA84 cells using a Seahorse XFp extracellular flux analyzer. The basal respiration rate and ATP production were higher in the KCL22-NTC cells compared to the Ptbp2-KO cells respectively (**Fig. 3A**), and lower in the LAMA84 vector control cells compared to the Ptbp2-OE-LAMA84 cells (**Supplementary Fig. 3A**). The spare respiratory capacity, defined as the difference between maximal and basal respiration, was found to be decreased by 55% in Ptbp2-KO-KCL22 cells. Furthermore, a similar decrease in the glycolysis rate was observed in the Ptbp2-KO-KCL22 compared to the KCL22-NTC cells (**Fig. 3B**) and decreased in the LAMA84-vector control compared to Ptbp2-OE-LAMA84 cells (**Supplementary Fig. 3B**). Ptbp2-KO-KCL22 and KU812 cells showed decreased intracellular ATP levels (**Fig. 3C and Supplementary Fig. 3C**). As expected, intracellular ATP levels increased in Ptbp2-OE-LAMA84 cells (**Supplementary Fig. 3D**). Thus, our data suggest that PTBP2 activates both oxidative phosphorylation (OXPHOS) and glycolysis.

**Figure 3.**
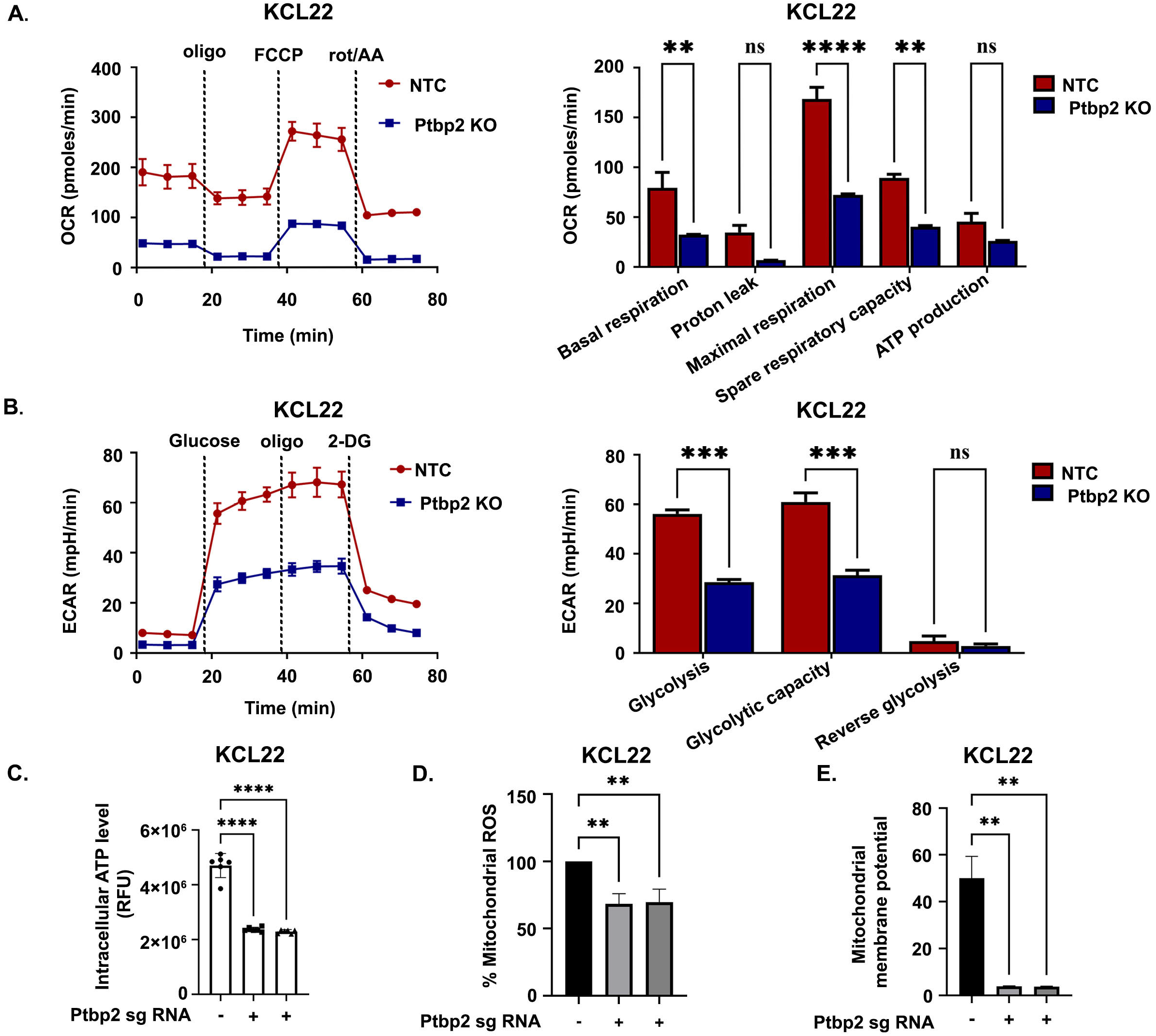
PTBP2 deficiency results in mitochondrial dysfunction. **A**. The Seahorse XFp Cell Mito Stress test was utilized, and the OCR measurement was examined in KCL22-NTC vs. Ptbp2-KO-KCL22 cells treated sequentially with oligomycin, FCCP, and rotenone (left panel). The outcome represents the mean and standard deviation of three separate experiments. Two-way ANOVA was used for statistical analysis, and the comparison test (right panel) **, p < 0.01, ****p < 0.0001, and ns-not significant. **B**. The glycolytic ability was measured in KCL22-NTC vs. Ptbp2-KO-KCL22 cells (left panel). The experiments were carried out using the Seahorse XFp Glycolysis Stress test, and the flow chart and bar graph depict the ECAR measurement (right panel). The results represent the mean and standard deviation of three separate experiments. Two-way ANOVA was implemented for statistical analysis, followed by a multiple comparison test ***p < 0.001 and ns-not significant. **C**. Intracellular ATP was assessed in KCL22-NTC vs. Ptbp2-KO-KCL22 cells. The ATP luminescence signals were normalized to the quantity of protein (n = 4), **** *p*< 0.0001. **D**. Flow cytometry assessed Mitochondrial superoxide distribution using MitoSOX dye and displayed by a bar graph in KCL22-NTC vs. Ptbp2-KO-KCL22 cells, n=2, ** *p*< 0.01. **E**. Flow cytometry measurement of MMP using JC-1 dye in KCL22-NTC vs. Ptbp2-KO-KCL22 cells, illustrated by a bar graph, ** *p*< 0.01, compared to the controls.

Respiration is essential for cellular proliferation and is correlated with biological aggressiveness. ROS-regulated signaling pathways are notably upregulated in different types of cancer, leading to cell proliferation, survival, and other factors contributing to cancer onset and progression (**Liou and Storz, 2010**). In addition to decreased ATP production and OXPHOS activity, lower levels of mitochondrial ROS were observed in Ptbp2-KO-KCL22, KU812 cells, and LAMA84-vector control cells (**Fig. 3D, Supplementary Fig. 3E and 3F**), including lower level of mitochondrial membrane potential in the Ptbp2-KO-KCL22 cells (**Fig. 3E**). Live cell imaging showed the exact change in the membrane potential (**Supplementary Fig. 3G**).

Mitochondrial fusion and fission play essential roles in mitochondrial morphology, and recent studies suggest that increased fusion correlates with increased Mitofusion-1 and 2 (MFN1 and MFN2) that boosts OXPHOS, leading to heightened cell proliferation (**Yao et al., 2019**). Thin and elongated mitochondria were observed under a confocal microscope in KCL22-NTC cells using MitoTracker staining, whereas, in PTBP2 KO cells, we observed dotted or fragmented mitochondria (**Fig. 4A, upper panel, and lower panel, respectively**). For further confirmation, we checked the expression of MFN1 and MFN2, which was seen to be reduced in the KO condition (**Fig. 4B**). We also examined the expression of dynamin-related protein 1 (DRP1), which drives constriction and fission during mitochondrial division. The Ptbp2-KO-KCL22 cells showed higher expression of DRP1 with respect to the KCL22-NTC cells (**Fig. 4B**). Thin and elongated mitochondria were also observed in the Ptbp2-OE-LAMA84 cells. In contrast, the vector control cells showed only dotted and fragmented mitochondria (**Supplementary Fig. 4A)**. Likewise, an increase in MFN1 and MFN2 expression and a decrease in DRP1 expression was observed in Ptbp2-OE-LAMA84 cells in comparison to vector control cells (**Supplementary Fig. 4B**). The transmission electron microscope has been very useful in studying the mitochondrial structure. Intact rod-shaped elongated mitochondria were observed in KCL22-NTC cells; however, damaged mitochondria were observed in Ptbp2-KO-KCL22 cells (**Fig. 4C**). Thus, PTBP2 promotes increased OXPHOS and mitochondrial fusion and thereby facilitates cell proliferation.

**Figure 4.**
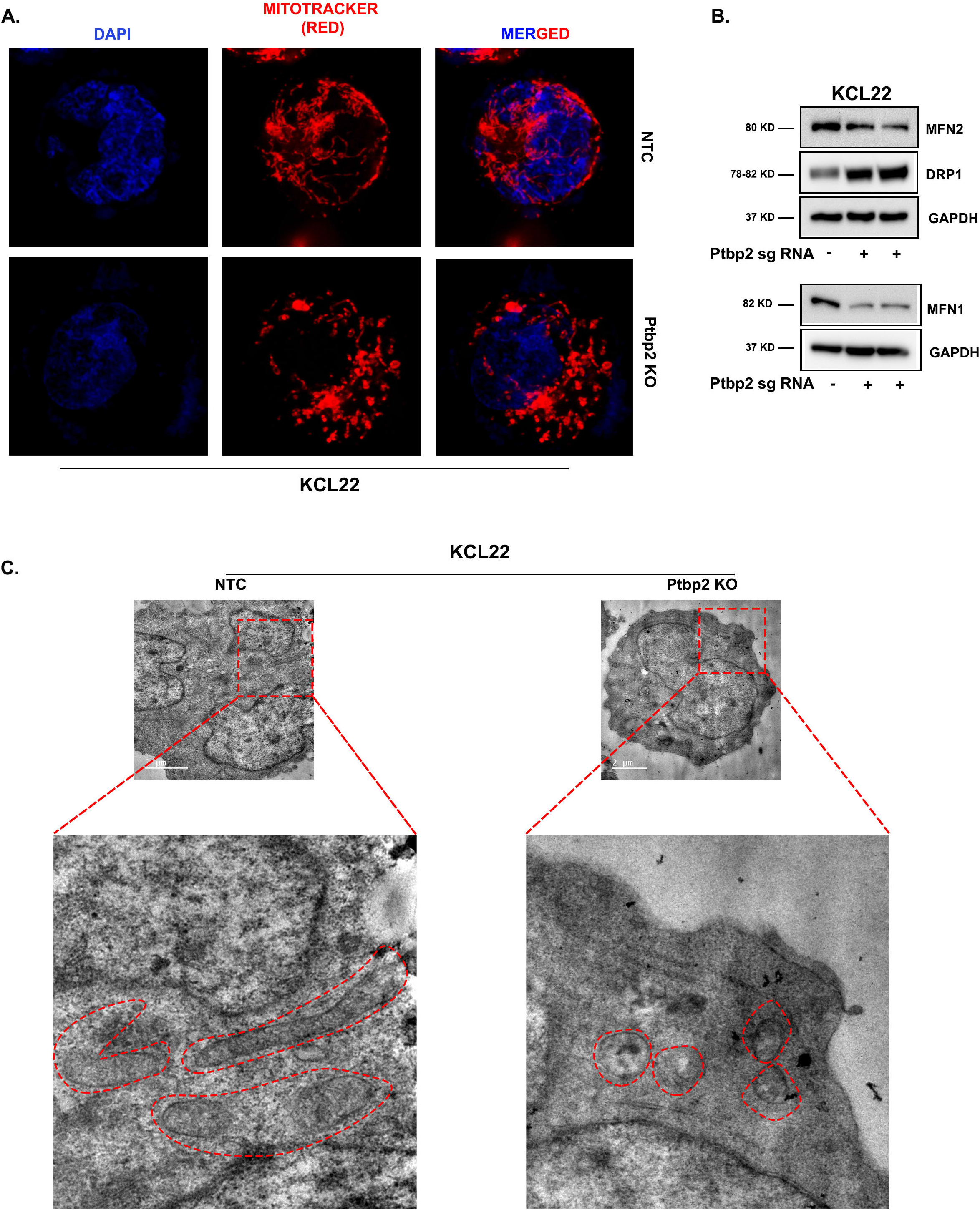
Diminution of PTBP2 affects mitochondrial morphology. **A**. Confocal microscopy image of KCL22-NTC vs. Ptbp22-KO-KCL22 cells using CMXros mitotracker red dye. **B**. Western blot analysis of MFN2 & DRP1 in control KCL22-NTC vs Ptbp2-KO-KCL22 cells. An equal amount of protein (30μg) was used for both cell lines, and GAPDH was used as an internal loading control (upper panel). Western blot analysis of MFN1 in control KCL22-NTC vs. Ptbp2-KO-KCL22 cells. An equal amount of protein (30μg) was used for both cell lines, and GAPDH was used as an internal loading control (lower panel). **C**. Representative TEM images of mitochondria in KCL22-NTC & Ptbp2-KO-KCL22 cells.

### The presence of PTBP2 promotes autophagy through BNIP3

Previous studies have suggested that BNIP3 can bind to Bcl-2 and release Beclin-1. This process promotes autophagy and inhibits apoptosis **(Zhang et al., 2008; Burton et al., 2009)**. In Ptbp2-KO-KCL22 and KU812 cells, the expression of Beclin-1 was decreased, whereas it was increased in LAMA84 cells where Ptbp2 was overexpressed (**Fig. 5A and 5B, 1**^**st**^ **panel**). The transformation of Microtubule-associated Protein 1 Light Chain 3B (LC3B) from its unconjugated state (LC3-I) to a phosphatidylethanolamine-conjugated state (LC3-II) plays a crucial role in the formation of autophagosomes **(Li et al., 2020)**. BNIP3 is primarily located on the mitochondria and has an LC3B interacting region (LIR) that facilitates the removal of mitochondria via autophagy **(Liu et al., 2022)**. ATG7 plays a role in converting LC3-I to LC3-II, and ATG12 plays a crucial role during autophagosome function. Both ATG7 and ATG12 were decreased in Ptbp2-KO-KCL22 and KU812 cells (**Fig. 5A, 2**^**nd**^ **and 3**^**rd**^ **panel**). Ptbp2-OE-LAMA84 cells showed upregulation of ATG7 and ATG12 (**Fig. 5B, 2**^**nd**^ **and 3**^**rd**^ **panel**). As BNIP3 is also involved in mitophagy, we examined the expression of mitophagy-related genes Optineurin and TOM20. Optineurin, an autophagy receptor in the parkin-mediated mitophagy pathway, was upregulated in KCL22-NTC, KU812-NTC, and Ptbp2-OE-LAMA84 cells compared to their counterparts (**Fig. 5A and 5B, 4**^**th**^ **panel**), whereas TOM20, whose abnormal expression is considered an indicator of mitophagy, was downregulated in KCL22-NTC, KU812-NTC, and Ptbp2-OE-LAMA84 cells (**Fig. 5A and 5B, 5**^**th**^ **panel)**. PTBP2 and GAPDH expression levels are also shown **(Fig. 5A, 6**^**th**^ **and 7**^**th**^ **panel, respectively)**. Furthermore, we overexpressed Bnip3 in Ptbp2-KO-KCL22 cells to determine the function of BNIP3 (**Fig. 5C**). To gauge the activity of autophagic flux, we used bafilomycin A1, an inhibitor of autophagosome-lysosome fusion, and studied the conversion of LC3-I to LC3-II. After treatment with Bafilomycin A1, KCL22-NTC cells showed higher expression of LC3-II than LC3-I, almost the same as in Ptbp2-KO-Bnip3-OE-KCL22 cells. The LC3-II/LC3-I ratio was quantified and shown in the figure (**Fig. 5D, 1**^**st**^ **panel**). Again, on treating cells with Bafilomycin A1, a partial increase in fragmented mitochondria in KCL22-NTC cells along with elongated ones was observed, but no difference in mitochondrial dynamics was observed in Ptbp2-KO-KCL22 cells (**Fig. 5E, upper and middle panel**). However, with the same condition, upon BNIP3 overexpression, in the Ptbp2-KO-KCL22 cells, the cells behaved as KCL22-NTC cells in terms of mitochondrial dynamics with a partial increase in elongated mitochondria, suggesting a possible role of BNIP3 in the process (**Fig. 5E, lower panel)**. On Bafilomycin A1 treatment, the number of small LC3 puncta was quantified. There was a significant reduction in the number of puncta in the PTBP2-KO-KCL22 cells with respect to the KCL22-NTC cells; however, upon overexpression of Bnip3 in the PTBP2-KO-KCL22 cells, the number of puncta was found to be increased (**Fig. 5F and Supplementary Fig. 5A**). P62 is an autophagy substrate used as an autophagy activity receptor. The lack of autophagy was clear by the accumulation of p62 only in the PTBP2-KO-KCL22 cells **(Fig. 5H, upper panel)** and not in KCL22-NTC and Ptbp2-KO-BNIP3-OE-KCL22 cells. A direct role of PTBP2 was observed when Ptbp2-OE-LAMA84 cells were used with respect to the vector control. The ratio of LC3-II/LC3-I was higher in Ptbp2-OE-LAMA84 cells than in the vector control cells (**Supplementary Fig. 5B, lane 4 with respect to lane 2**). These results suggested that PTBP2 plays a role in the autophagy of CML cells. Ablation of Mfn2 led to impaired autophagic degradation and suppressed cell proliferation **(Ding et al., 2015; Chen and Dorn, 2013)**. To understand whether this is mediated through BNIP3, we used a scrambled siRNA and BNIP3-specific siRNA to knock down the expression of BNIP3 in KCL22 cells. Upon successful knockdown of BNIP3, MFN2 was found to be reduced, demonstrating that BNIP3 works through MFN2 (**Fig. 5I, 1**^**st**^ **panel**). In the same lysate, we checked for the expression of Beclin-1 & ATG7, which was found to be reduced. Thus, our data suggest that BNIP3 promotes autophagy through the Beclin-I pathway **(Fig. 5I, 2**^**nd**^ **and 3**^**rd**^ **panel)**.

**Figure 5.**
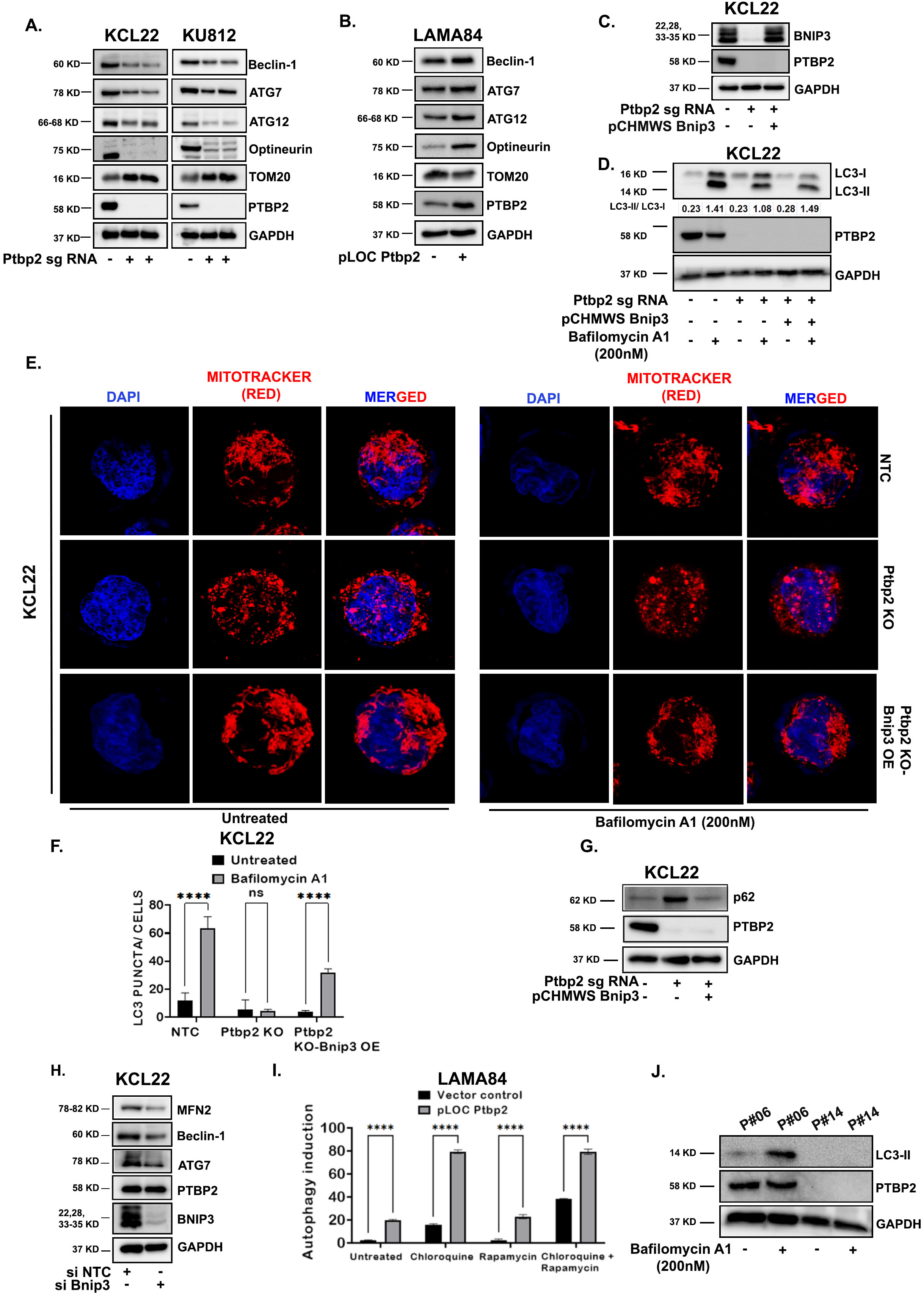
Regulation of autophagy by PTBP2 in CML cells. **A**. Western blot analysis of Beclin-1, ATG7, ATG12, Optineurin, TOM20, and PTBP2 in KCL22-NTC vs. Ptbp2-KO-KCL22 cells (left panel) & in KU812-NTC vs Ptbp2-KO-KU812 cells (right panel). An equal amount of protein (30μg) was loaded & GAPDH was used as an internal loading control. **B**. Representative immunoblotting analysis of Beclin-1, ATG7, ATG12, Optineurin, TOM20, and PTBP2 in vector control LAMA84 vs Ptbp2-OE-LAMA84 cells. **C**. Western blot analysis was performed for BNIP3 in KCL22-NTC, Ptbp2-KO-KCL22, and Ptbp2-KO-Bnip3 OE-KCL22 cells by taking an equal amount of protein. GAPDH was considered a loading control. **D**. KCL22-NTC, Ptbp2-KO-KCL22, Ptbp2-KO-Bnip3-OE-KCL22 cells were treated with bafilomycin A1 (200nM) for 18h, lysed and immunoblotted for LC3B with an equal amount of protein (30ug). A western blot was also performed using the Ptbp2 antibody. GAPDH was used as a loading control. Densitometry quantification is shown in the figure. **E**. Confocal microscopic image of KCL22-NTC, Ptbp2-KO-KCL22 & Ptbp2-KO-Bnip3-OE-KCL22 cells using CMXros mitotracker red dye before and after treatment with Bafilomycin A1 (200nM) for 18h. **F**. Representative bar diagram of LC3 puncta in KCL22-NTC, Ptbp2-KO & Ptbp2-KO-Bnip3-OE-KCL22 cells treated with Bafilomycin A1 (200nM) for 18h. *** *p*< 0.001. **G**. Western blot analysis was performed for p62 & PTBP2 in KCL22-NTC, Ptbp2-KO-KCL22, and Ptbp2-KO-Bnip3-OE-KCL22 cells by taking an equal amount of protein. GAPDH was considered a loading control. **H**. KCL22 cells were transfected with scrambled siRNA and Bnip3 siRNA, and after 48h, cells were lysed, and immunoblotting was performed for DRP1, MFN2, Beclin-1, ATG7, PTBP2, & BNIP3 by taking an equal amount of protein. GAPDH was used as a loading control. **I**. Flow cytometry-based profiling of autophagy in LAMA84 vector control vs. Ptbp2-OE-LAMA84 cells was done. Cells were treated or untreated with 0.5μM rapamycin, 10μM chloroquine (CQ), or both for 20h. After staining with CYTO-ID green detection reagent for 30 mins in the dark, cells were washed and analyzed by flow cytometry. Results were analyzed and represented in a bar diagram. *** *p*< 0.001. **J**. Immunoblotting for LC3B was performed in two patient samples treated with Bafilomycin A1 (200nM) for 18h.

The Cyto-ID autophagy detection kit measures autophagic vacuoles and monitors autophagic flux in lysosomally inhibited live cells. Chloroquine (CQ), an antimalarial drug, inhibits autophagy by preventing the fusion of autophagosomes with lysosomes and slowing lysosomal acidification. Ptbp2-OE-LAMA84 cells, when treated with CQ or combined with rapamycin, showed higher autophagic flux than the vector control cells (**Fig. 5J, bars 4 and 8, with respect to bars 3 and 7**). The autophagic flux was reduced when only rapamycin was used without chloroquine (**Fig. 5J, bars 5 & 6**).

We looked for the LC3B turnover in CML patient samples for clinical relevance. The patient sample expressing PTBP2 showed LC3-II expression after treatment with Bafilomycin A1, whereas in the sample that did not express PTBP2, even after Bafilomycin A1 treatment, LC3-II was found to be absent **(Fig. 5K)**. Our observation suggests that PTBP2-mediated stabilization of Bnip3 promotes autophagy and enhances cell survival.

### KCL22 cells promote tumorigenesis in athymic nude mice

To investigate the effect of PTBP2 *in vivo*, exponentially growing KCL22-NTC, Ptbp2-KO-KCL22, and Ptbp2-KO-Bnip3-OE-KCL22 cells were subcutaneously injected into the flanks of athymic nude mice. From the second week, tumor volume was measured using a digital caliper. The animal study plan is shown in **Fig. 6A**. We observed large tumors in mice injected with KCL22-NTC cells compared to those injected with Ptbp2-KO-KCL22 cells (**Fig. 6B, upper and middle panel**). However, the introduction of Ptbp2-KO-Bnip3-OE-KCL22 cells resulted in tumors of size between the wild type and Ptbp2 KO cells **(Fig. 6B, lower panel)**. Tumor weight and volume were higher in KCL22-NTC cells than in Ptbp2-KO-KCL22 and Ptbp2-KO-Bnip3-OE-KCL22 cells (**Fig. 6C and 6D, respectively**). In addition, a significant decrease in PTBP2, BNIP3, and Ki-67 expression was observed in the Ptbp2-KO-KCL22 cells with respect to the NTC, as confirmed by IHC (**Fig. 6E, 1**^**st**^ **and 2**^**nd**^ **column**). However, upon overexpression of BNIP3 in the Ptbp2-KO-KCL22 cells, expression of Ki-67 and Beclin-1 was found to be increased (**Fig. 6E, 3**^**rd**^ **column**).

**Figure 6.**
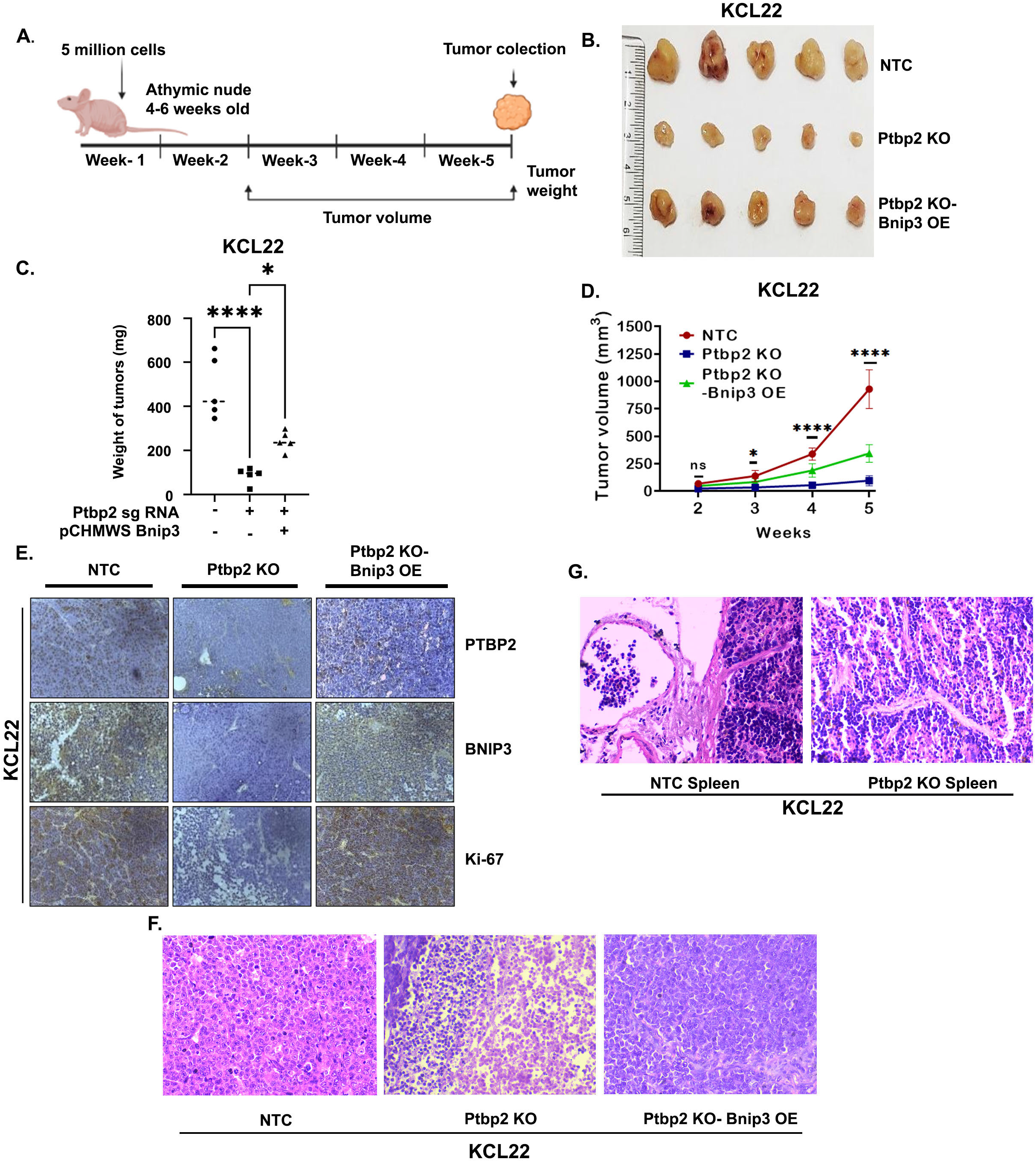
PTBP2 promotes tumor growth. **A**. Plan of work for the *in vivo* study. Five million respective cells were injected subcutaneously in 4-6 weeks-old female athymic nude mice. After five weeks, the tumor was collected and processed for IHC. In between that, tumor volume was measured twice per week. **B**. Representative tumor images are represented in the mice injected with KCL22-NTC, Ptbp2-KO-KCL22, and Ptbp2-KO-Bnip3-OE-KCL22 cells. The experiment was performed two times, with 5 animals in each group, and the Vernier caliper was used to measure the volume. **C**. Graphical representation of the tumor weight of each group of mice and the average weight of tumors were represented by scatter plot. t-test was performed to calculate the statistical significance between those groups. *, *p*< 0.05, ****, *p*< 0.0001. **D**. The rate of tumor progression of KCL22-NTC, Ptbp2-KO-KCL22 & Ptbp2-KO-Bnip3-OE-KCL22 cells is represented by the tumor volume calculated each week. Data is represented by a line graph with an error bar. *, *p*< 0.05, ****, *p*< 0.0001 **E**. Representative IHC images showing the PTBP2, BNIP3, and Ki-67 expression pattern in KCL22-NTC, Ptbp2-KO-KCL22, and Ptbp2-KO-Bnip3-OE-KCL22 group tumor. **F**. The tumor tissue was stained with H&E and visualized at x400. **G**. The spleen tissue was stained with H&E and visualized at x400.

In tissue samples derived from the tumors of NTC mice, apparently healthy and actively dividing cells were observed upon H&E staining. The cells were arranged in diffuse sheets supported by well-vascularized fibrous tissue septa. Cellular pleomorphism, a characteristic feature of malignancy, was more evident in mice with KCL22-NTC tumors than in tissue samples derived from tumors of Ptbp2-KO-KCL22 **(Fig. 6F)**. KCL22-NTC cells showed large blasts with scant to moderate cytoplasm. The nuclei were vesicular, round, oval, or sometimes folded, with finely stippled chromatin, prominent nucleoli, and high mitotic activity. Some cells were very large, with a high nuclear-cytoplasmic ratio and prominent nucleoli. In the Ptbp2-KO-KCL22 group, large areas of necrosis were observed with small islands of proliferating neoplastic cells in between. Individual cell necrosis was evident, characterized by highly eosinophilic cytoplasm and loss of nuclei or karyolysis. The neoplastic cells in the active areas were mainly a uniform population of immature cells with some intermediate forms and without much pleomorphism. In the Ptbp2-KO-Bnip3-OE-KCL22 group, the cells were primarily consistent with a lower mitotic rate than in the NTC group but higher than that in the KO group. The migration of KCL22-NTC cells through the subcapsular sinus and trabeculae into the spleen was observed. However, no migration of cells into the spleen was observed in mice injected with Ptbp2-KO-KCL22 **(Fig. 6G)**. Thus, we conclude that high levels of PTBP2 may contribute to developing a more aggressive form of the disease.

## Discussion

RBPs, including Musashi2 (Msi2), heterogeneous nuclear ribonucleoprotein H1, hnRNPA1, and others, have been implicated in CML (**Kharas et al., 2010; Zhang et al., 2014; Li et al., 2020; Wang M. et al., 2020; Liu, et al., 2021**). Msi2 expression was significantly increased, followed by Ptbp2 expression, in blast crisis samples compared to other RBPs (**Nandagopalan et al., 2019**). PTBP2 has been associated with oncogenic RNA splicing in various cancers, such as glioblastoma, osteosarcoma, and colorectal cancer **(Xu and Hecht, 2011; Nowak et al., 2011; Ji et al., 2014; Yang et al., 2014; Tang et al., 2023**). Knockdown of both PTBP1 and PTBP2 slowed glioma cell proliferation (**Cheung et al., 2009)**, and PTBP2 null mice died shortly after birth and exhibited misregulation of alternative splicing in genes involved in cytoskeletal remodeling and cell proliferation **(Boutz et al., 2007; Licatalosi et al., 2012**). We found that the knockout of Ptbp2 in several CML and AML cells reduced proliferation and long-term colony formation ability. We demonstrated that PTBP2 could increase glycolysis and OXPHOS, leading to a more significant proliferation of cells through increased ATP production and possibly elevated macromolecular precursors. **Yao et al., 2019** recently reported that glycolysis and mitochondrial respiration support the proliferation of oncogene-transformed cells, which aligns with our observation. MFN1 and MFN2 are proteins found in the outer membrane of mitochondria that primarily function as a mitochondrial fusion protein. Mitochondrial biogenesis, as assessed by fusion, was increased in CML cells, which was counteracted when Ptbp2 was knocked out. The elongated shape of the mitochondria in KCL22-NTC cells and the presence of MFN1/MFN2 suggest that excessive fusion of mitochondria drives cell proliferation. DRP1 is the primary regulator of mitochondrial fission. In Ptbp2-KO-KCL22 cells, the existence of DRP1 alongside dotted mitochondria suggests excess fission. Increased mitochondrial fission in Ptbp2 conditional KO (cKO) spermatids was shown due to increased DRP1. This leads to differences in the number and shape of mitochondria between WT and Ptbp2 cKO spermatids **(Hannigan et al., 2018**).

Mitochondria, the primary source of ROS in most mammalian cells, are typically higher in cancer cells and function as crucial signaling molecules regulating autophagy and tumor development. Several reports indicate that inhibition of autophagy may be a promising approach for the treatment of BCR-ABL-mediated leukemia **(Karvela et al., 2016; Helgason et al., 2011)**. In this study, we demonstrate that PTBP2 binds to a subset of mRNAs with a very high affinity, of which BNIP3 is one, and was found to be stabilized by PTBP2. Despite its pro-death activity, BNIP3 expression in cancer often predicts an aggressive disease. Upregulation of BNIP3 characterizes cancer cell subpopulations with increased fitness and proliferation (**Zhu et al., 2022)**. We also observed a strong correlation between Ptbp2 and Bnip3 expression in CML patients. Induction of autophagy through BNIP3 in pre-invasive breast cancers provides tumor cells with extra nutrients and promotes tumor progression **(Zhang et al., 2022)**. Mechanistically, the LC3-II/LC3-I ratio decreased when PTBP2 was knocked out and increased when BNIP3 was overexpressed in the knockout cells. It was observed that the expression of Beclin-1 and ATG7 decreased in KO cells and also in KCL22-NTC (siRNA BNIP3) cells. We previously showed the antagonistic expression of PTBP1 and PTBP2 in CML cells (**Nandagopalan et al., 2019**). Thus, consistent with our findings, the knockdown of PTBP1 (which increases the level of PTBP2) has been reported to lead to autophagy in colorectal and bladder cancer (**Takai, et al., 2017)**. Also, the knockdown of PTBP1 caused the transition of LC3-I to LC3-II in breast and bladder cancer cells, while overexpression of PTBP1 reduced the transition of LC3-I to LC3-II (**Takai, et al., 2017; Wang et al., 2018)**. Fission is followed by selective fusion that segregates dysfunctional mitochondria and permits their removal by autophagy **(Twig et al., 2008)**. These reports and our data strongly suggest that the PTBP2-BNIP3 axis can positively regulate the proliferation of CML cells through the combinatorial action of mitochondrial fusion and the transition of LC3-I to LC3-II, thereby promoting autophagy. The formation of large tumors with microscopic features of cellular pleomorphism and anaplasia and increased Ki67 expression in mice injected with KCL22-NTC cells in comparison to Ptbp2-KO-KCL22 xenografts provide convincing evidence that the presence of PTBP2 simultaneously promotes proliferation, leading to aggressive tumor formation. This suggests that PTBP2 acts as an oncogene in CML and that PTBP2-mediated induction of autophagy through BNIP3 may provide CML cells with extra nutrients and promote further tumor progression. Based on our current and past data, it is worth proposing PTBP2 as a potential biomarker and therapeutic target for controlling CML progression. In addition, PTBP2 protected cells from the clinically approved BCR-ABL1 inhibitor imatinib, as Ptbp2 KO cells that failed to show autophagy were more sensitive to imatinib-mediated cell death. Thus, targeting PTBP2-BNIP3-mediated autophagy by genetic ablation of PTBP2 or pharmacological inhibition of PTBP2 can reduce disease severity and enhance imatinib sensitivity in CML cells.

## Materials and Methods

### Cell lines and patient samples

The CML (KCL22, KU812, K562, KYO1, & LAMA84), AML (TF1, HEL, HNT34, & F36P) cell lines and HEK293T cells were cultured in RPMI medium 1640 (Gibco, Cat. #31800014) +10% fetal bovine serum (FBS) (Gibco, Cat. #10270106) and Dulbecco’s modified Eagle medium (Gibco, Cat. #12100046) respectively, with no antibiotic supplements. Blood samples were collected from CML patients, and the study was approved by the institutional human ethical committee (16/HEC/12).

### Generation of stable Ptbp2 KO cell lines

The knockout cell lines were produced by transducing specific sgRNAs for Ptbp2 exon 1 (Horizon, USA, Cat. #VSGH11936-247759039, Ptbp2 sgRNA-1) and exon 2 (Horizon, USA, Cat. #VSGH11936-2430473, Ptbp2 sgRNA-2) in KCL22, KU812, and TF1 cell lines. The Edit-R All-in-one Lentiviral sgRNA non-targeting control (NTC) viral particles (Horizon, USA, Cat. #VSGC11954) were used to obtain the control cells. The clonal selection was performed in those transduced cells, and RT-qPCR and western blotting confirmed PTBP2 knockout.

### Generation of stable cell lines overexpressing PTBP2 and BNIP3

The study used lentivirus-based precision lentiORF PTBP2 w/stop codon to overexpress Ptbp2 (Horizon, USA, Cat. #OHS5899-202618163) in LAMA84 cells. For control cells, lentivirus-based precision LentiORF positive control (pLOC) (Horizon, USA Cat. #OHS5833) was transduced into LAMA84 cells and selected with blasticidin. For overexpression of Bnip3, the coding region of Bnip3 was cloned into a lentivirus vector; lentivirus particles were produced by transfecting target plasmid into HEK293T cells, sorting GFP-positive cells and transducing the same in PTBP2 knockout cells. The expression of PTBP2 was checked by using RT-qPCR (supplementary methods).

### Western blotting

Cell protein lysates were obtained using radioimmunoprecipitation assay buffer (RIPA) (Cell Signalling Technology (CST), Cat. #9806). Western blot analysis and the antibodies used are mentioned in the supplementary methods.

### RNA immunoprecipitation

According to the manufacturer’s instruction, the EZ-Magma NuclearTM RIP (cross-linked) nuclear RNA-binding protein immunoprecipitation kit (Sigma-Aldrich, Cat. #17-10521) was used. Briefly, 30×10^6^ cells were taken and fixed with 0.75% formaldehyde. After fixation, cells were spun down and re-suspended in a buffer comprising protease and RNase inhibitors. The suspension was then incubated on ice for 15 mins, followed by centrifugation at 4°C. Lastly, the pellet was re-suspended in RIP cross-linked lysis buffer (RCB) and incubated on ice, followed by sonication to prepare the lysate. The prepared lysate was split into two fractions, and antibodies (PTBP2 in one fraction & IgG in another) were added, followed by protein G beads (Millipore, Cat. #16-266) to the immuno-complex for 2h. The co-precipitated RNAs were isolated and sequenced (Supplementary methods).

### Cell proliferation assay

5x10^5^ cells were seeded in a T25 flask in 7ml of RPMI-1640+10% FBS for each group, and the culture was maintained for four days. Each day, the cells were counted independently three times using a Trypan blue solution (Sigma-Aldrich, Cat. #T8154) for each group for the number of viable cells, and their average was considered. A clonogenic assay was conducted, as mentioned in the supplementary methods.

### Measurements of OCR and ECAR

The oxygen consumption rate (OCR) and Extracellular Acidification Rate (ECAR) were measured using a Seahorse XFp extracellular flux analyzer (Agilent Technologies, USA). Before the experiment, the XFp-mini plate was coated with 100μl per well of poly-L-lysine and left for at least 45 min at 37°C. Each well was washed with Milli-Q water and dried for at least 1h. 1.25×10^5^ KCL22-NTC and Ptbp2 KO cells were seeded in each well in 180μl of XF RPMI Assay Medium (Seahorse Bioscience, Cat. #103576–000) supplemented either with 2mM L-glutamine (Cat. #P0480100) for ECAR measurement or 1mM Sodium-pyruvate (Gibco, Cat. #11360-070), 10mM glucose (Gibco, Cat. #A24940-01), 2mM L-glutamine for OCR analysis, as per the manufacturer’s instructions. The plate was centrifuged and left to equilibrate for 60 min in a CO_2_-free incubator before being transferred to an XFp extracellular flux analyzer (Agilent technology, USA). Mitochondrial membrane potential (MMP), cell viability assay, and ROS production assay were conducted as mentioned in the supplementary methods.

### Autophagy detection assay

Per the manufacturer’s instruction, autophagy detection was performed using CYTO-ID green detection reagent 2 (Enzo Life Sciences, Cat. #ENZKIT175). Briefly, cells were treated with chloroquine or rapamycin for 20h, then centrifuged, rinsed, and suspended in an assay buffer with 5% FBS. Diluted CYTO-ID green was added to each sample, incubated, and analyzed using flow cytometry.

### Animal study

All animal protocols were performed with the approval of the ethics committee of the Institute of Life Sciences (ILS/IAEC-130-AH/AUG-18). Cells were injected subcutaneously into the lower flank of 4-6-week-old nude mice. Tumor growth was monitored, and volumes were measured every three days from the second week. Five weeks after injection, mice were sacrificed, and tumor weight was measured and analyzed. Tumors were fixed in 10% neutral buffered formalin for the H&E staining and IHC examination. All studies involved five mice per group. Immunohistochemistry was conducted per the standard protocol and is mentioned in the supplementary methods.

## Supporting information

All supplementary data

## Acknowledgment

BB and SL are supported by a fellowship from the Council of Scientific and Industrial Research (CSIR), India. SC was supported by a fellowship from the University Grants Commission (UGC), India. The pCMV PTBP2 plasmid was kindly provided by Dr. Miriam Llorian (University of Cambridge, Department of Biochemistry, Cambridge, UK). The authors would like to acknowledge the help of Mr. Bhabani Sahoo of the Confocal facility, Mr. Paritosh Nath of the Flow Cytometry facility, Dr. Sarita Jena, and Mr. Biswajit Patra of the Animal House facility.

## Author contribution

BB, SL, and SC* designed the experiments. BB and SL performed the experiments. SC^¥^ performed RT-qPCR. SIS performed the H&E staining and interpretation. SM, SB, and GB provided clinical inputs. SC* arranged for funding, supervised the experiments, interpreted the data, and wrote the manuscript. All authors reviewed and approved the final version of the manuscript. SC^¥^: Sayantan Chanda; SC*: Soumen Chakraborty.

## Data Access Statement

Research data supporting this publication are available on request.

## Ethical approval and consent to participate

All animal protocols were approved by the Institute of Life Sciences (ILS/IAEC-130-AH/AUG-18) Animal Ethics Committee. The Institutional Human Ethical Committee approved the study (16/HEC/12), and peripheral blood samples were collected from patients after obtaining written informed consent. All procedures involving human participants were performed according to the ethical standards of the institutional research committee and with the 1964 Helsinki Declaration and its later amendments or comparable ethical standards.

