## Supplementary material for "PTBP2 promotes cell survival and autophagy in Chronic Myeloid Leukemia by stabilizing BNIP3": All supplementary data

### Supplementary figure 1

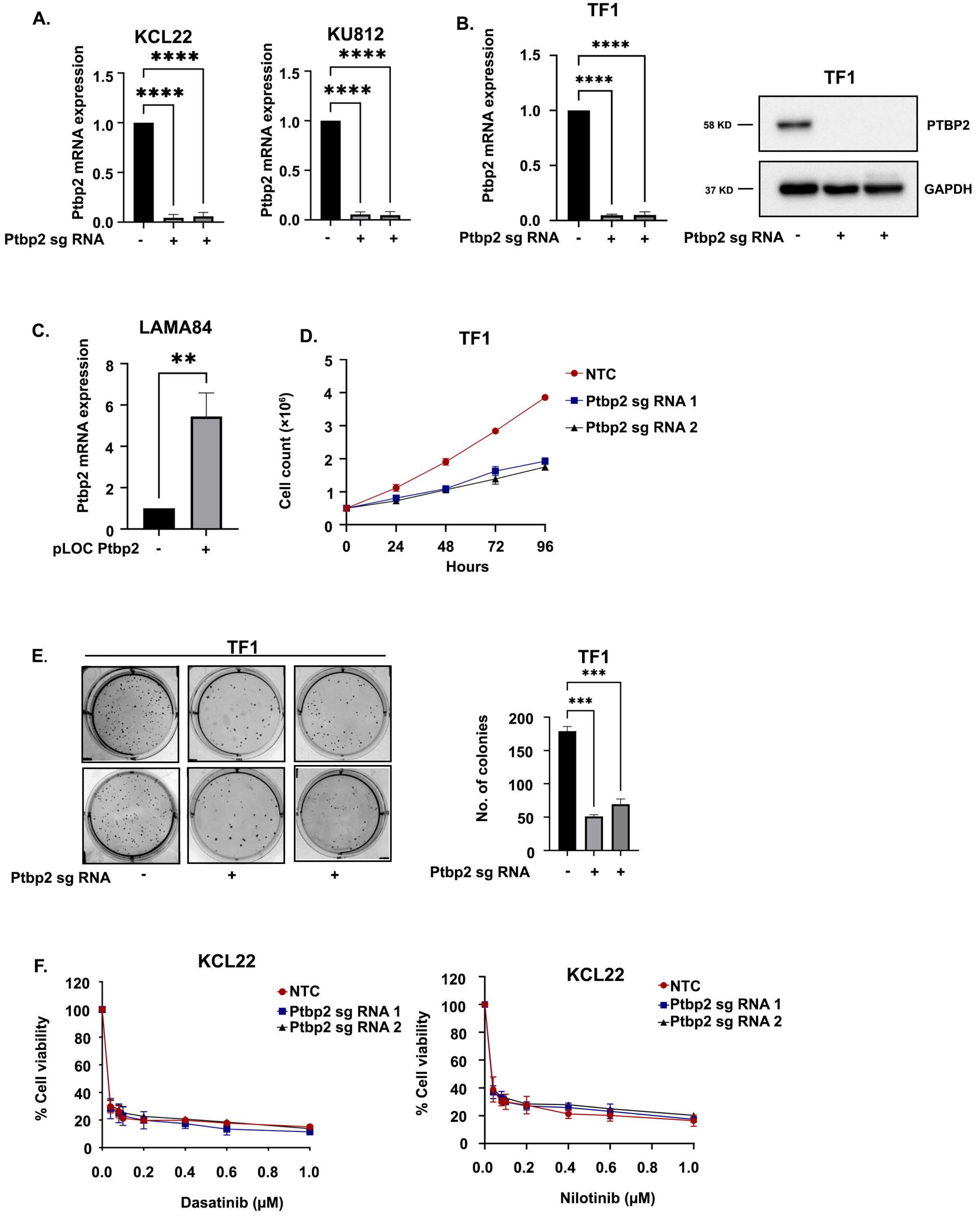

### Supplementary figure 2

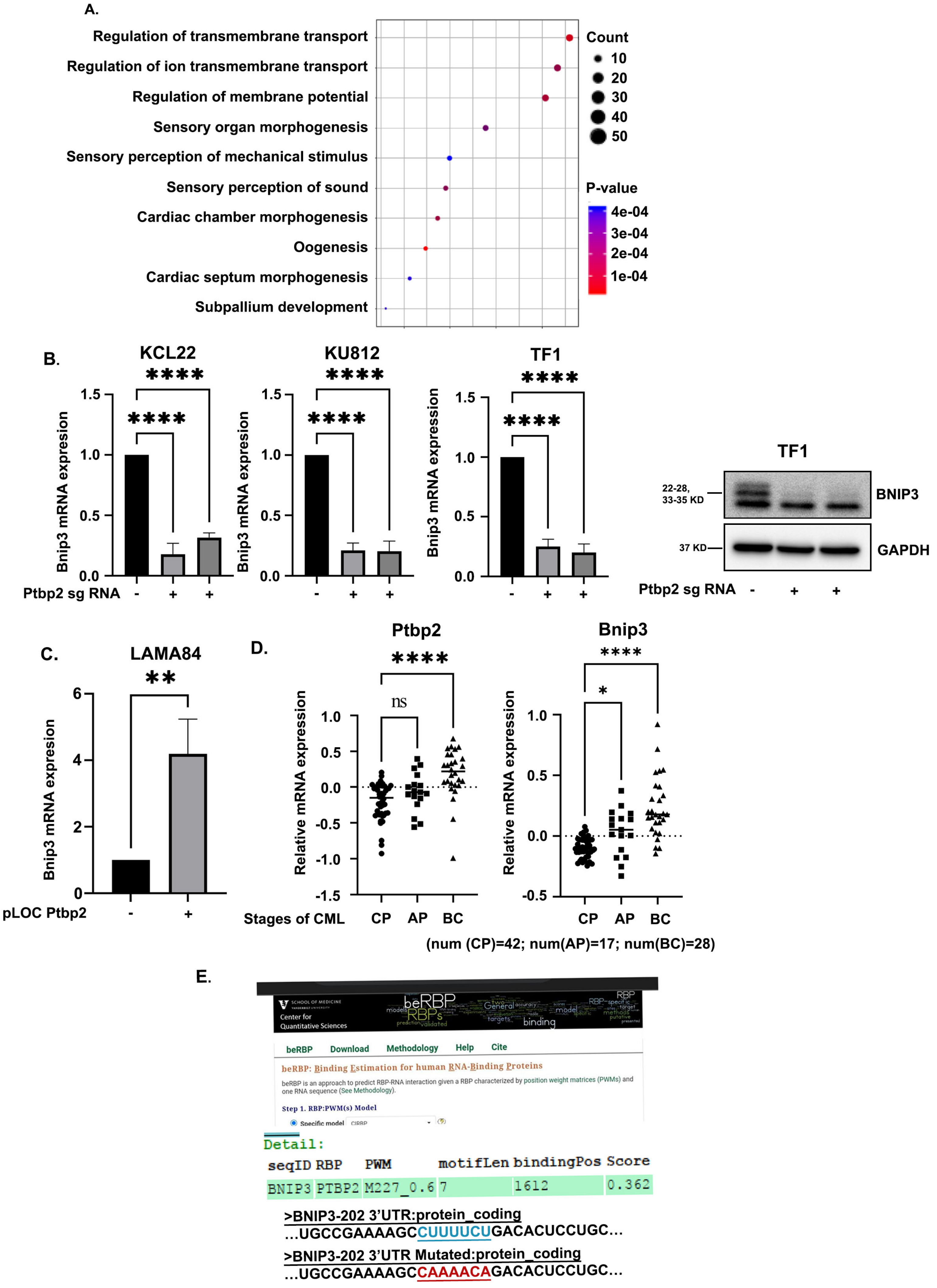

### Supplementary figure 3

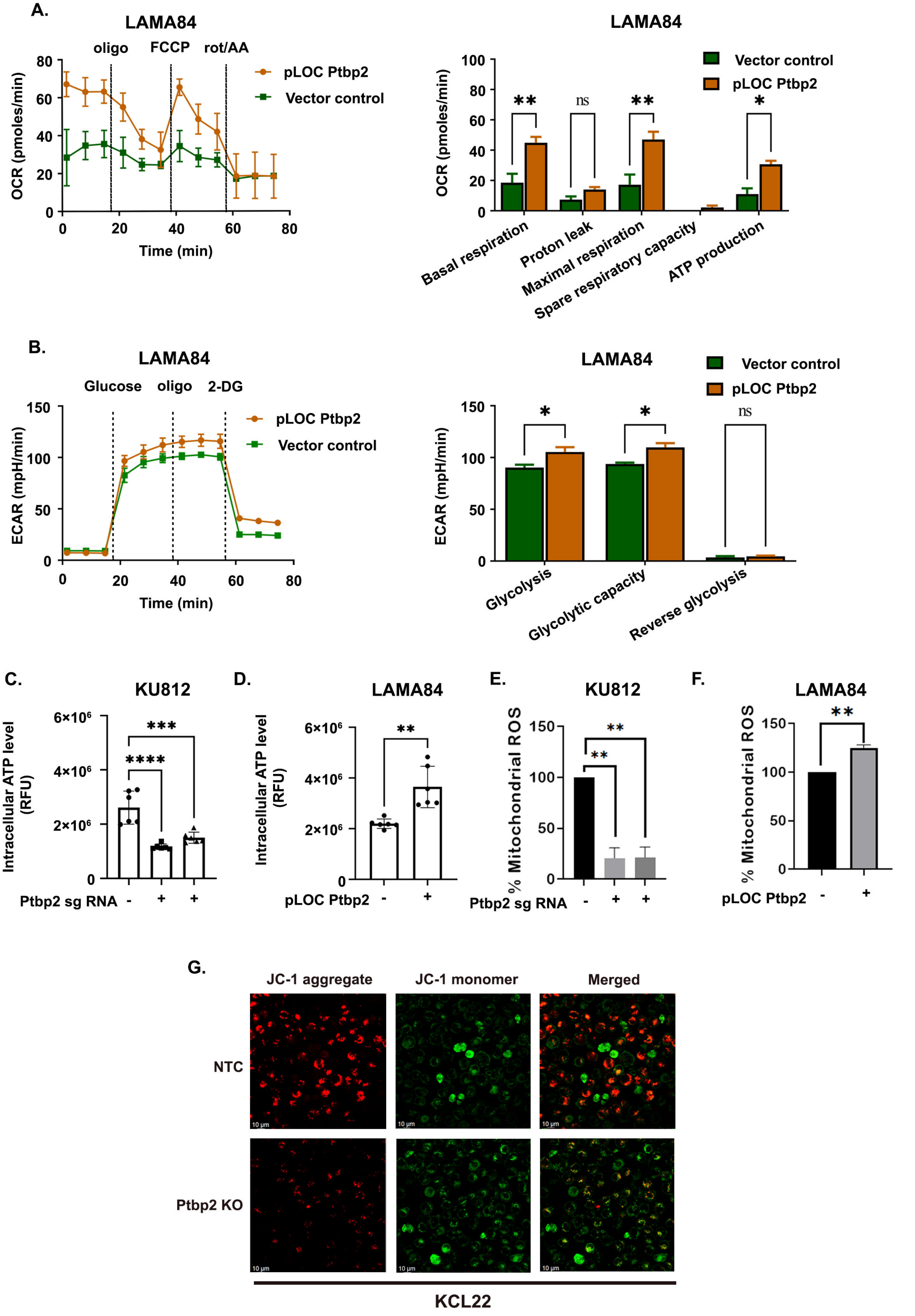

### Supplementary figure 4

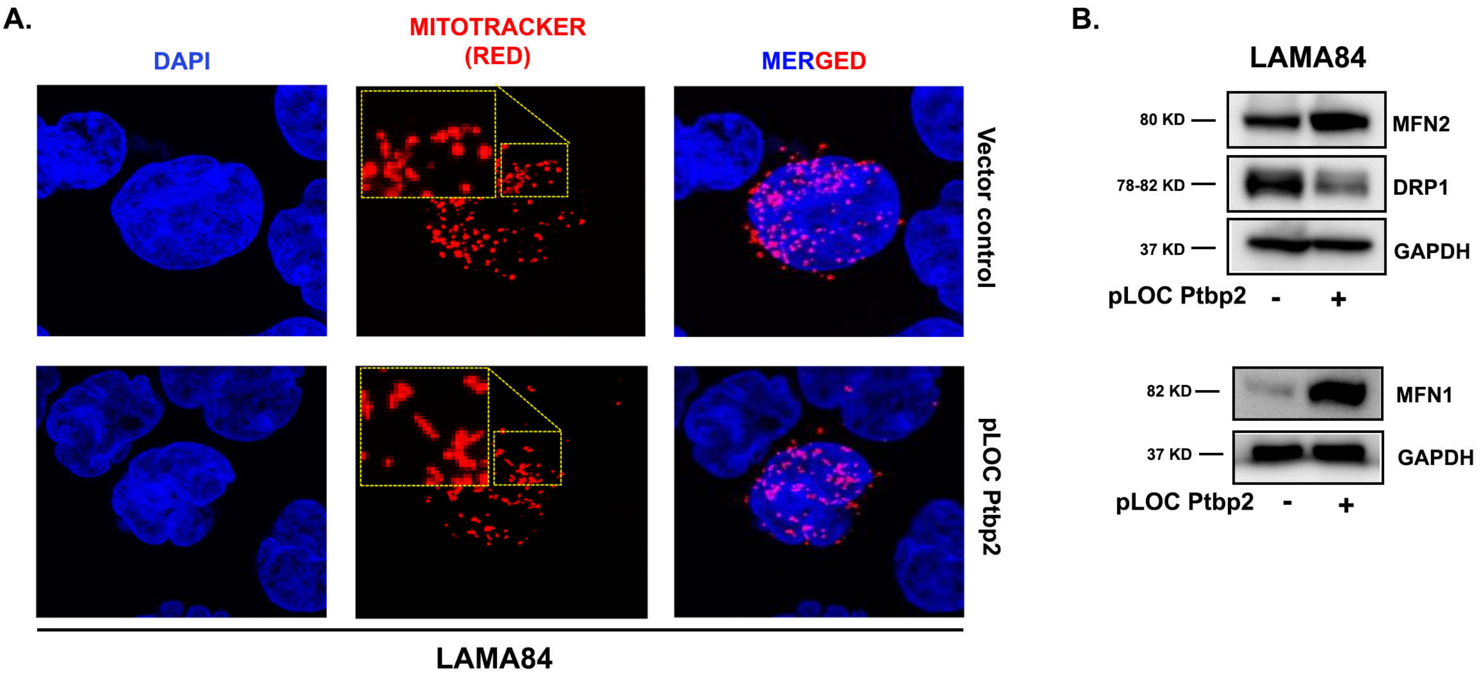

### Supplementary figure 5

**A.**

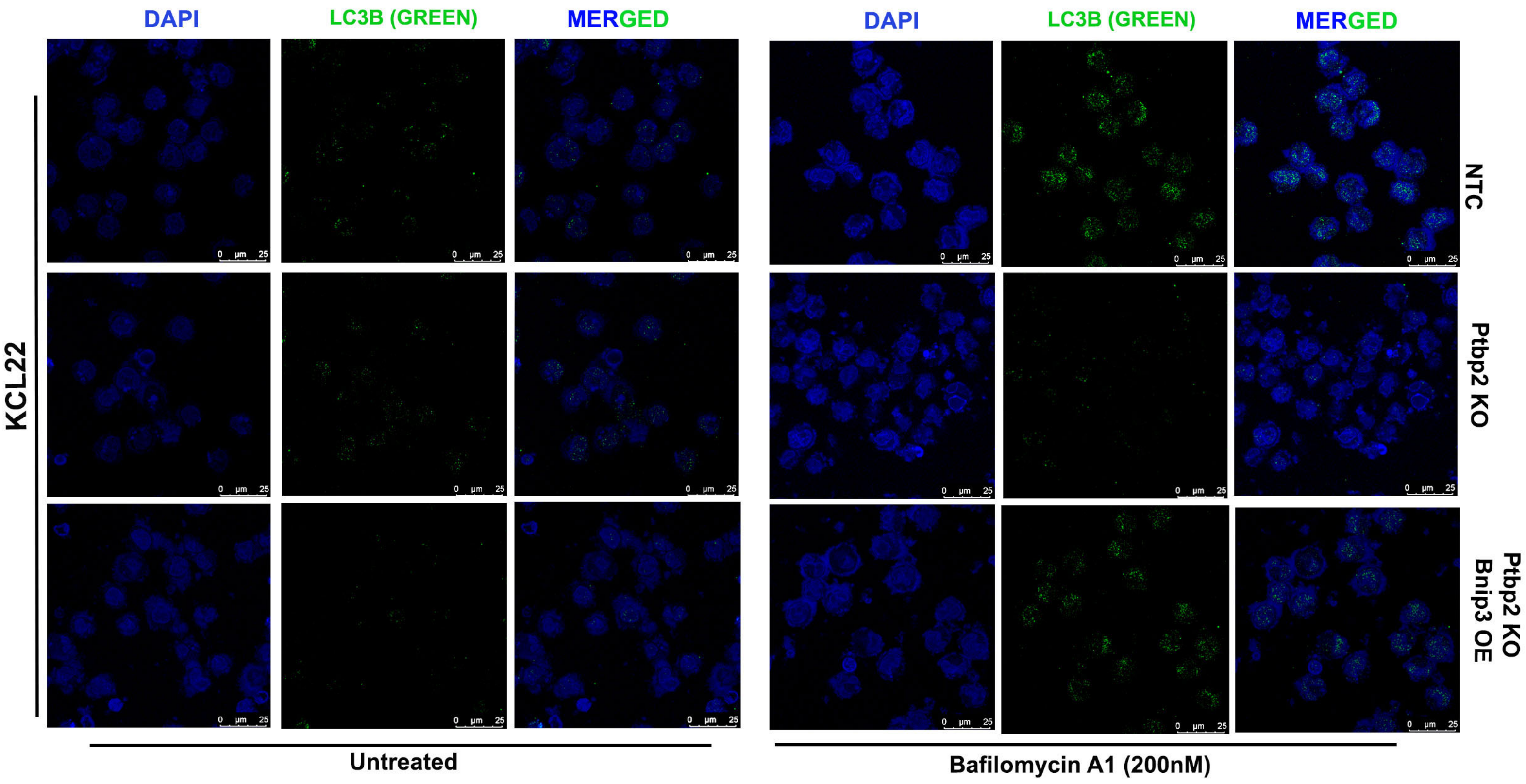


**B.**

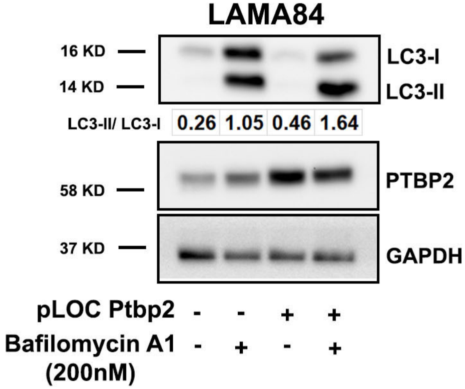

#### Supplementary Figure legends

**Supplementary Figure 1: PTBP2 protein expression in different cell types and phenotypic assessment. A**. mRNA expression of Ptbp2 using RT-qPCR in the KCL22-NTC, KU812-NTC, Ptbp2-KO-KCL22, and Ptbp2-KO-KU812 cells. **B.** The mRNA expression of Ptbp2 using RT-qPCR in the TF1-NTC and Ptbp2-KO-TF1 cells. The protein expression of Ptbp2 was checked through immunoblotting in TF1-NTC and Ptbp2-KO-TF1 cells. **C**. mRNA expression of Ptbp2 using RT-qPCR in the LAMA84 vector control and Ptbp2-OE-LAMA84 cells. **D.** TF1-NTC and Ptbp2-KO-TF1 cell proliferation rates were checked using a trypan blue exclusion test for cell viability. The proliferation rates at 24, 48, 72, and 96h are represented in the line graph. Each point represents the mean and standard deviation of independent triplicates. **E.** The soft agar colony formation assay was performed with TF1-NTC and Ptbp2-KO-TF1 cells. The colony numbers are represented in the bar graph below the respective figures. The bar diagram shows the mean value and corresponding standard deviation for the represented data; $n=3$. ***$P < 0.001$. **F.** Representative graph of % cell viability in KCL22-NTC & Ptbp2-KO-KCL22 cells using variable doses of Dasatinib & Nilotinib.

**Supplementary Figure 2: PTBP2 binds and regulates BNIP3. A**. Gene ontology study of common PTBP2 bound transcripts using Cluster profiler Bioconductor package (*$p<0.05$). The dot color represents the GO term's significance level, and the dot's size represents the number of genes enriched for the term. Color scale and size scale are given as legends. **B**. RT-qPCR analyzed the relative expression of BNIP3 in KCL22-NTC, KU812-NTC, Ptbp2-KO-KCL22, and Ptbp2-KO-KU812 cells. RT-qPCR was used to analyze BNIP3 mRNA expression in TF1-NTC and Ptbp2-KO-TF1 cells. Whole-cell lysates of TF1-NTC and Ptbp2-KO-TF1 cells were collected, immunoblotting was performed with BNIP3 antibody, and GAPDH was used as an internal loading control. **C.** RT-qPCR analyzed the relative expression of Bnip3 in LAMA84 vector control and Ptbp2-OE-LAMA84 cells. **D.** Relative expression of Ptbp2 (left panel) & Bnip3 (right panel) in different phases of CML obtained from patient samples in Radich Oncomine data set (GSE4170). CP (n=42), AP (n=17), and BC (n=28). **E.** Schematic representation of the binding site of PTBP2 in the 3'UTR region of Bnip3 was detected by beRBP software. The binding region is underlined (top), and the site was mutated by site-directed mutagenesis, as represented below.

**Supplementary Figure 3: PTBP2 deficiency results in mitochondrial dysfunction. A**. Mitochondrial stress tests of the vector control LAMA84 vs. Ptbp2-OE-LAMA84 cells. The Seahorse XFp Cell Mito Stress test was utilized, and the OCR measurement was examined (left panel). The outcomes represent the mean and standard deviation of three separate experiments. Two-way ANOVA was used for statistical analysis and the comparison test (right panel). **B**. The glycolytic capability was measured in vector control LAMA84 vs. Ptbp2-OE-LAMA84 cells (left panel). The experiments were carried out using the Seahorse XFp Glycolysis Stress test, and the flow chart and bar graph depict the ECAR measurement (right panel). The results represent the mean and standard deviation of three separate experiments. Two-way ANOVA was implemented for statistical analysis**. C**. Intracellular ATP was assessed in KU812-NTC vs. Ptbp2-KO-KU812 cells. The ATP luminescence signals were normalized to the quantity of protein (n = 4). **D.** Intracellular ATP was assessed in vector control LAMA84 vs Ptbp2-OE-LAMA84 cells. The ATP luminescence signals were normalized to the quantity of protein (n=4). **E**. Mitochondrial superoxide distribution was assessed using mitoSOX dye by flow cytometry and displayed by a bar graph in KU812-NTC vs. Ptbp2-KO-KU812 cells. **F**. Mitochondrial superoxide distribution was assessed using mitoSOX dye by flow cytometry and displayed by a bar graph in vector control LAMA84 vs. Ptbp2-OE-LAMA84 cells. **G**. Flow cytometry measurements of MMP using JC-1 dye in KCL22-NTC vs. Ptbp2-KO-KCL22 cells illustrated by live cell imaging.

**Supplementary Figure 4: Diminution of PTBP2 affects mitochondrial morphology. A.** Confocal microscopic image of vector control LAMA84 vs. Ptbp2-OE-LAMA84 cells using CMXros mitotracker red dye. **B**. Western blot analysis of MFN2 & DRP1 in vector control LAMA84 vs Ptbp2-OE-LAMA84 cells. Protein was loaded in equal amounts (30µg). GAPDH was used as an internal loading control (Upper panel). Western blot analysis of MFN1 in vector control LAMA84 vs Ptbp2-OE-LAMA84 cells. Protein was loaded in equal amounts (30µg). GAPDH was used as an internal loading control.

**Supplementary Figure 5: Regulation of autophagy in CML cells. A.** Confocal microscope image of KCL22-NTC, Ptbp2-KO-KCL22, and Ptbp2-KO-Bnip3-OE-KCL22 cells using LC3B primary antibody & 488 fluorophore secondary antibody after treatment with bafilomycin A1 for 18h. **B.** Western blot for LC3B was performed in vector control vs. Ptbp2-OE-LAMA84 cells treated with bafilomycin A1 (200nM) for 18h, and GAPDH was taken as a loading control. The normalized densitometry ratio of LC3-II/LC3-I was taken, and values are represented in the figure.

#### Supplementary Materials and Methods

##### RNA isolation, cDNA preparation, and RT-qPCR

RNA was isolated from the cell lines and patient samples using the TRIzol method (Ambion, Cat. #15596018) and quantified using NanoDrop 2000/2000c (Thermo Fisher Scientific, USA). The first-strand cDNA synthesis kit (Thermo Fisher Scientific, Cat. #K1612) was synthesized using reverse transcriptase with 2µg of RNA. RT-qPCR was performed using the GoTaq qPCR master mix (Promega, Cat. #A6002). GAPDH was used as a loading control. Primer sequences used in this study are listed in Supplementary Table 1.

##### RNA Immunoprecipitation Sequencing (RIP-Seq)

RNA sequencing libraries were prepared using Illumina-compatible NEBNext® Ultra™ II Directional RNA Library Prep Kit (New England BioLabs, MA, USA) following the manufacturer's instructions. Briefly, 1ng of total immunoprecipitated RNA was taken for fragmentation and priming. Fragmented and primed RNA was further subjected to first-strand synthesis followed by second-strand synthesis. The double-stranded cDNA was purified using HighPrep Beads (Magbio, Cat # AC-60050). Purified cDNA was end-repaired, adenylated, and ligated to Illumina multiplex barcode adapters as per NEBNext® Ultra™ II Directional RNA Library Prep protocol followed by second strand excision using USER enzyme at 37˚C for 15 min. Adapter ligated cDNA was purified using HighPrep Beads and was subjected to 16 cycles for Indexing- (98˚C for 30 sec, cycling (98˚C for 10sec, 65˚C for 75sec) and 65˚C for 5min) to enrich the adapter-ligated fragments. The final product (sequencing library) was purified using HighPrep Beads and a library quality control check. An Illumina-compatible sequencing library was quantified using a Qubit fluorometer (Thermo Fisher Scientific, MA, USA), and its fragment size distribution was analyzed using an Agilent 2100 Bioanalyzer. After sequencing, the raw data generated was checked for quality using FastQC. Reads were preprocessed to remove the adapter sequences and remove the low-quality bases (<q30). Pre-processing of the data is done with CutAdapt. HISAT2, a spliced aligner, was used to align the high-quality data with the default parameters of the reference genome. RIPSeeker was used for RIP-seq analysis, which uses machine learning on an alignment file to first model peak enrichment within the IP sample while considering unique hits. Then, it reassigns multiple hits to their most likely location based on posterior probability, according to a "rich get richer" model. Peak calling is then performed again on the unique and disambiguated multi-hits. It produced a list of peaks and their corresponding candidate-bound transcripts for each valid entry.

##### Western blotting

We used 30-50µg of cell lysate with DTT sample loading buffer, resolved them in a sodium dodecyl sulfate-polyacrylamide gel, and transferred them onto an Immobilon-P polyvinylidene fluoride membrane (Merck, Cat. #IPVH00010). Blots were probed with the indicated primary antibodies and horseradish peroxidase-conjugated secondary antibodies. For blot development, Amersham™ ECL™ Prime Western blotting Detection Reagent (GE Healthcare, Cat. #RPN2232) on a Chemidoc™ MP Imaging System (Bio-Rad, USA). The primary antibodies used in this study were anti-GAPDH (Cat. #2118S) and PTBP2 (Cat. #15719) and BNIP3 (Cat. #44060S), and LC3B (Cat. #3868S), ATG7 (Cat. #8558T), ATG12 (Cat. #4180), Beclin-1 (CST, Cat. #3495T), and MFN2 (Cat. #11925), DRP1 (CST, Cat. #8570), TOM20 (CST, Cat. #42406), and Optineurin (Cat. #79028) and normal Rabbit IgG (CST, Cat. #2729S), and secondary mouse antibody (Cat. #7076), and secondary rabbit antibody (Cat. #7074). All the antibodies were purchased from Cell Signaling Technology, USA.

**Clonogenic assay**

1.2% of low melting agarose gel was prepared in Milli-Q water, which was autoclaved and then mixed with 2x RPMI-1640 growth media in a 1:1 ratio to prepare the final 0.6% gel (before mixing, both the solutions were brought to 37°C). 2 ml/well of the prepared gel solution was plated in a low-attachment six-well plate as the lower layer. Similarly, the upper layer was prepared at a final concentration of 0.35% and contained 500 cells per well. 250µl of feeding media (RPMI) was administered twice weekly. After 21 days, the colonies were counted and analyzed using the ImageJ software (NIH, Bethesda, MD, USA).

**Confocal microscopy**

KCL22 NTC and Ptbp2 KO cells, LAMA84 vector control, and Ptbp2 OE cells were treated with 50nM MitoTracker® Red CMXRos (Invitrogen, Cat. #M-7512) for 20 mins at 37°C. The cells were washed and placed on a coverslip coated with poly-L-lysine (Cat. #P4707). After fixing the cells with 4% paraformaldehyde (PFA) (Sigma-Aldrich, Cat. #P6148) for 20 min, they were permeabilized with 0.1% Triton X-100 (Sigma-Aldrich, Cat. #X100) for 10 min. The coverslips were then mounted on slides using a mounting medium containing DAPI (Thermo Fisher Scientific, Cat. #D1306) for nuclear staining and visualized using a confocal microscope at 63x (Leica, Germany).

**Measurement of MMP**

To measure the mitochondrial membrane potential (MMP), $1\times10^5$ KCL22 NTC and Ptbp2 KO cells were stained with JC-1 dye, a cationic fluorescent carbocyanine dye (Sigma-Aldrich, Cat. #T4069) at 37 °C in the dark for 30 min. Cells were washed with 1x PBS, centrifuged (400×g for 5 min), and resuspended in PBS. Flow cytometry (BD Biosciences, USA) and live cell imaging (Discover, Zeiss) detected fluorescence.

**MTT assay**

For the cell viability assay, $1 \times 10^4$ cells/well were seeded in 96-well plates and treated with different concentrations of imatinib. After 48h, 10µl of 0.5 mg/mL MTT prepared in $dH_2O$ (Sigma-Aldrich, Cat. #M5655) was added to each well and incubated for 4h at 37°C to form formazan crystals, which were dissolved in DMSO. The absorbance was measured at 570 nm using a VICTOR Nivo$^{TM}$ multimode reader (PerkinElmer, USA).

**ROS production assay**

Reactive Oxygen Species (ROS) production assays were performed in NTC, Ptbp2 KO, and Ptbp2 OE cells. Cells ($1\times10^5$) were placed in a 1.5 ml tube and washed with 1x PBS. Single-cell suspensions were made using 200µl of accutase treatment for 5 min at 37°C. An extra 500µl of media was added to neutralize the accutase reaction and centrifuged at 300×g for 5 min at 4°C; then, the medium was discarded. Cells were washed once with 1x PBS and incubated with 300µl of 5µM MITOSOX (Thermo Fisher Scientific, Cat. #M36008) (5µM) for 30 min at 37°C in the dark. Then 200µl of 1x PBS was added, and ROS production was quantified using flow cytometry.

**Immunohistochemistry**

Formalin-fixed and paraffin-embedded 4-5µm thick sections were deparaffinized, rehydrated, and quenched in 3% $H_2O_2$ for 20 min. The cells were washed with phosphate-buffered saline (PBS) (Gibco, Cat. #14190-094) and blocked in PBS containing 2.5% horse serum for 1hr at RT. The tissues were incubated overnight at 4°C with the required dilution of primary antibodies. The slides were washed three times in 1x PBS and incubated with biotinylated secondary antibodies (Vector Laboratories, Cat. #30082) for 45 min, followed by incubation with avidin-biotin complex (Vector Laboratories, Cat. #PK-7200) for 30 minutes at RT. Immunoreactivity was determined by using diaminobenzidine as the final chromogen. Finally, the sections were counterstained with Delafield hematoxylin (Hi-Media, India, Cat. #S014), dehydrated using increasing concentrations of ethanol, cleared in xylene, and mounted with a DPX mounting medium. Each slide was marked with one of the sections as a negative control, on which the primary antibody was omitted. Images were obtained using a Leica DM500 microscope (Leica, Germany). The antibodies used for the assay were as follows: Ki 67 (CST, Cat. #9027S) (1:200) and BNIP3 (CST, Cat. #44060S) (1:100), PTBP2 (Millipore, Cat. #ABE431) (1:100).

**Luciferase reporter assay**

The luciferase gene was cloned into the pCDNA3.1 vector's HindIII, BamHI site after being removed from the pGL3 vector. Concurrently, PCR was used to amplify BNIP3-3' UTR, which contains the anticipated PTBP2 binding site, and TA cloning was used to clone it into the PGMT vector. After that, EcoRI was used to digest the PGMT vector to separate the BNIP3-3' UTR site along with the EcoRI restriction site and ligate it into a pCDNA3.1 vector (digested with EcoRI). The binding site was mutated by modified primers using site-directed mutagenesis (The primer sequence is mentioned in Table 1). Using Fugene HD transfection reagent (Promega, Cat.# E2311), HEK293T cells were transfected with the reporter constructs pCDNA3.1 luciferase BNIP3, pCDNA3.1 luciferase mutated BNIP3, pCMV mammalian expression vector containing PTBP2 gene, pCMV vector negative control, and Renilla luciferase gene (pRL-TK). Cells were treated and lysed 48 hours after transfection using the PLB buffer included in the kit (Promega, Cat# Dual luciferase reporter assay system, E1910). After centrifuging the sample, the luciferase activity was measured using a luminometer (GLOMAX 20/20, Promega, USA). The results were normalized against the Renilla luciferase activity as an internal control. All measurements were done in triplicates, and the assay was repeated thrice.

**Transmission emission microscopy (TEM)**

Cells were washed thrice with 0.1M sodium cacodylate buffer (all the further washing steps performed with this buffer) and fixed in 2.5% glutaraldehyde prepared in 0.1M sodium cacodylate buffer at 4°C overnight. After three washes, the cells were post-fixed in 1% osmium tetraoxide for 2h at RT, followed by a wash. Next, the cells were stained with 1% uranyl acetate counterstain for 1h. After washing, cells were subjected to serial dehydration with 30%, 50%, 70%, 90%, and 100% ethanol (10 min at each dilution). The 100% ethanol dehydration step was repeated, followed by 100% propylene oxide. After dehydration, cells were resuspended in propylene oxide and epons resin mixture (Ted Pella, Cat. #183004221) (1:1, 1:2, 1:4 ratio respectively) each for 1h (no washing step required). Lastly, cells were resuspended in epons resin only and left with the cap open at RT overnight for the evaporation of leftover propylene oxide. The next day, cells were resuspended in epons resin mixed with hardener, and the cell mixture was transferred to fresh 0.5ml tubes and centrifuged to pellet down the cells to the bottom of the tube. The tubes were then undisturbed at 65°C for 48h for polymerization and hardening. Once the blocks were ready, ultra-thin sections (70nM) were cut using an ultramicrotome and collected on a copper grid, followed by staining with 2% uranyl acetate for 20 min and then with 1% lead citrate for 5 more min with washing step-in between with autoclaved MilliQ water (3 deeps in water). The sections were allowed to dry before capturing any images.

**Actinomycin D chase assay**

$3\times10^6$ KCL22-NTC, Ptbp2-KO-KCL22 cells were seeded a day before the experiment. The following day, cells were treated with Actinomycin D (Sigma Aldrich, Cat. #A4262) 5µg/ml to shut off transcription. Total RNA was isolated at different time points 1h, 2h, 4h, 6h, 8h, and Bnip3 mRNA levels were quantified by qRT-PCR. GAPDH was used as a loading control. Decay curves were plotted using GraphPad Prism software.

**siRNA transfection**

According to the manufacturer's protocol, the siRNA transfection was performed in KCL22 cells ($5\times10^5$) by Lipofectamine™ RNAiMAX Transfection Reagent (Thermo Fisher Scientific, Cat. #13778075). The transfected cells were confirmed for knockdown by immunoblotting after 48h of transfection. Bnip3 siRNA (Cat, #4392420), negative control siRNA (Cat. #AM4611) was obtained from Thermo Fisher Scientific.

**Live dead assay**

The cell viability cells were observed in a high-content analyzer using fluorescent live/dead dye (Thermo Fisher Scientific, Cat No# L3224). In brief, cells were treated with a sublethal dose of imatinib for 48h. After treatment, $1\times10^4$ cells were seeded in a black, flat-bottomed 96-well plate (Thermo ScientificTM Nunc) and stained with Calcein AM and Ethidium homodimer solution prepared in PBS. After 30min of staining at 37°C, plates were centrifuged at 1500 rpm, and photographs in individual wells were taken with the help of Cell Insight CX7 High Content Screening (HCS) Platform (20 fields per well using 10X objective lens) based on intensity. The green fluorescence indicates the live cells, and the red fluorescence indicates dead cells. Two fluorescent channels (excitation wavelengths −488 nm and 561 nm) were used to acquire green and red fluorescence. Image analysis was performed using the HCS Studio software.

**Statistical analysis**

The study used ImageJ software for protein band quantification from immunoblotting, GraphPad Prism 9.0 for data analysis, and a two-tailed unpaired student t-test for group comparisons. OCR and ECAR data were analyzed using WAVE 2.6.3. Statistical significance was determined at $p < 0.05$, ***, $p < 0.001$, and $p < 0.001$, and all experiments were repeated at least twice with biological repeats. All data are represented as mean ± SEM.

**Supplementary Table:** List of primers

| | | |
|---|---|---|
| 1 | PTBP2-F | TCTGAGTCCTTTGGCCATTC |
| 2 | PTBP2-R | GGTAAACAGACTTTGGGGCGTA |
| 3 | GAPDH-F | ACCCACTCCTCCACCTTTG |
| 4 | GAPDH-R | CTCTTGTGCTCTTGCTGGG |
| 5 | BNIP3 qRT-F | CTGCCGCCTTGATGTTTCACC |
| 6 | BNIP3 qRT-R | CTGAGAGATGACTGCTTCCAAGGT |
| 7 | KLHL28 qRT-F | CGCTGCCCGTGTGATTACTG |
| 8 | KLHL28 qRT-R | CAGAGTTCGTGATGTTGGCGA |
| 9 | UBXN2B qRT-F | TTCAGCGCCTTGTTCATGGT |
| 10 | UBXN2B qRT-R | TTGCCCTTCTCCACTAAAAGCC |
| 11 | TMEM106B qRT-F | TGGTTACTAAAGTTGGGGTGGC |
| 12 | TMEM106B qRT-R | AACAGCATCACATGCTAGAGACA |
| 13 | EMB qRT-F | TGAGGGAGCAGTCTCCACGA |
| 14 | EMB qRT-R | GTAAAAGGCGAATCTGGGGCAC |

| | | |
|---|---|---|
| 15 | ERO1L qRT-F | TCCTGAGCGCTACACTGGTTA |
| 16 | ERO1L qRT-R | GTTCTCTTCACTTGTCCCTTGACC |
| 17 | FAM63B qRT-F | CCATGGGTGGTTAGTAGACCCT |
| 18 | FAM63B qRT-R | GCTCAGCTACAAAGCCTTCACT |
| 19 | NAMPT qRT-F | AGTGACTTAAGCAACGGAGCG |
| 20 | NAMPT qRT-R | CTGCCGCAGGATTCATCTCG |
| 21 | PSMD11 qRT-F | AGCAAGGCTAAAGCAGCTCG |
| 22 | PSMD11 qRT-R | TGACTTGGCCCATTCGATGC |
| 23 | MCCC2 qRT-F | AGGAGGAGCCTATGTGCCTG |
| 24 | MCCC2 qRT-R | CCCAGTTGCCGCTTTAACCA |
| 25 | PAPOLA qRT-F | GCGTTGTGTGTTGCACCAAG |
| 26 | PAPOLA qRT-R | ACGAATGCCTCTTCAACAGCTC |
| 27 | ZNF681 qRT-F | GTGGCAATGCCTGGACACTAT |
| 28 | ZNF681 qRT-R | GTTCCAGACAGGTGATCAGGTC |
| 29 | C6orf25 qRT-F | CCCATGGGTCCGTGTATCCC |
| 30 | C6orf25 qRT-R | TCTGGGGCTCGGTTTTCACA |
| 31 | DKK2 qRT-F | CCACCGAGATGGCATGTGCT |
| 32 | DKK2 qRT-R | GCCGAGTACCATCCAGAGCC |
| 33 | ACTR2 qRT-F | TGCCATCACGGTTGGAACGA |
| 34 | ACTR2 qRT-R | TGCGGGGTGGGTCTTCAATG |
| 35 | UBE2V2 qRT-F | AAAATGTAGAGGACACAGCTGGGAA |
| 36 | UBE2V2 qRT-F | ACAGTTTTGATGGCAGGCTACAGT |
| 37 | PTP4A1 qRT-F | AAGCAGGCTCGACACTGAGAC |
| 38 | PTP4A1 qRT-R | AGGAATCATGCTGCAAACCACG |
| 39 | BBX qRT-F | TGAGTGTTTTCAGTCTCTCCGGG |
| 40 | BBX qRT-R | GTGTGTAGCTAGCCCTCAGCG |
| 41 | CCN1 qRT-F | ACGTAGCTTGCCCCGTCAAA |
| 42 | CCN1 qRT-R | TGGAGAAGCAAGCAAAAGCAACAG |
| 43 | DENR qRT-F | CGACCGCCTGGGTTTGTGAA |
| 44 | DENR qRT-R | GGACTCGAAGTGGGTAATCGGC |
| 45 | TWSG1 qRT-F | TGGGACGAGTGCTGTGACTG |
| 46 | TWSG1 qRT-R | CCTTCTGTGAGTGCCCGGAA |
| 47 | BNIP3-3’UTR PTBP2 binding site-F | TGCCGAAAAGCCTTTTCTGACACTCCTGCA |
| 48 | BNIP3-3’UTR mutant PTBP2 binding site-F | TGCCGAAAAGCCAAAACAGACACTCCTGCA |
| 49 | BNIP3-3’UTR-R | CTACCAGGGCAAGATGGTGT |
